# Unifying Gene Regulatory Network Inference and Sequence-to-Expression Modeling via Bilinear Sparse Regression for De Novo TF–DNA Interaction Discovery

**DOI:** 10.64898/2026.11.26.769492

**Authors:** Jordi Alonso Esteve, Dick de Ridder, Aalt D.J. van Dijk

## Abstract

Inferring transcription factor (TF)-DNA interactions from transcriptomic data is key to deciphering the genomic regulatory code. Existing computational approaches generally focus either on identifying trans-regulatory dynamics via gene regulatory network (GRN) inference or cis-regulatory sequence logic via sequence-to-expression (S2E) modeling. This requires fitting separate models across individual genes or conditions, which obscures shared regulatory structures and fails to explicitly capture cis-trans interactions. Here, we introduce BRIDGE (Bilinear Regulatory Inference from DNA and Gene Expression), a computationally efficient framework that unifies GRN inference and S2E modeling into a single, convex optimization objective. By explicitly modeling gene expression as a function of the interaction between promoter *k*-mers and TF abundances, BRIDGE guarantees a unique global minimimum while enabling *de novo* discovery of TF-specific binding motifs (TFBMs). Without the use of prior motif libraries, BRIDGE reconstructs high-fidelity TFBMs matching experimentally validated ones. Hence, we establish a new conceptual framework for linking TF abundance to cis-regulatory sequence, providing a foundation for future approaches to build upon.

## Introduction

Gene expression in eukaryotes is regulated by the coordinated action of trans-regulatory factors, including transcription factors (TFs), microRNAs or chromatin remodelers, and cis-regulatory elements (CREs), such as promoters and enhancers. The dynamic interplay between these components gives rise to a rich diversity of gene expression programs that underlie cellular identity and function. Understanding this interplay is fundamental to deciphering the regulatory code of the genome.

Decoding this system requires uncovering the rules of regulatory specificity. This is a challenging task: a typical eukaryotic genome encodes hundreds to thousands of distinct TFs that can bind thousands to hundreds of thousands of potential cis-regulatory targets to execute highly precise, reliable expression programs. Regulatory genomics assays such as ChIP-seq, SELEX or DAP-seq (Bartlett et al., 2017) have been instrumental in characterizing the relationship between TFs and CREs, providing experimental evidence of TF binding and its sequence preferences. However, these approaches provide only a partial view of the regulatory landscape. TF binding is highly context-dependent and can be influenced by cellular state, chromatin accessibility, cooperative interactions with other TFs or epigenetic modifications (Slattery et al., 2014). Moreover, the large number of TFs, regulatory elements, and experimental conditions makes systematic characterization of regulatory interactions difficult to scale across the genome (O’Malley et al., 2016).

In contrast, RNA-seq offers a comparatively inexpensive and highly scalable readout of gene expression, enabling regulatory programs to be studied across large numbers of cell types, tissues, conditions and perturbations. Although RNA-seq does not directly measure TF–CRE interactions, it acts as an indirect sensor of regulatory activity, capturing the downstream gene expression consequences of those interactions. Owing to it, uncovering TF-CRE interactions from RNA-seq data became a classical problem in computational biology (Bussemaker et al., 2001; Balwierz et al., 2014). Here, we use transcription factor binding motifs (TFBMs) to refer to DNA sequence motifs that represent the sequence preferences associated with TF binding.

Purely data-driven approaches for the analysis of RNA-seq data in the pursuit of prediction and/or understanding of gene expression can be broadly categorized into two perspectives, which emphasize two complementary aspects of gene regulation: sequence-to-expression (S2E) modeling and gene regulatory network (GRN) inference. In both cases, the TF–CRE interactions that substantiate transcription are more or less the implicit target, as we show below.

Early approaches to motif discovery from gene expression data followed a two-step strategy. First, genes were clustered according to their expression profiles across different conditions. The promoters of genes within each cluster were then searched for overrepresented sequence motifs using tools such as MEME (Bailey et al., 1994; Tompa et al., 2005; Aerts et al., 2024; Tavazoie et al., 1999). However, similarity in expression profiles, for example as measured by correlation, does not necessarily imply co-regulation, nor does it guarantee that genes are regulated through the same set of motifs. Hence, more recent S2E approaches formulate motif inference as a regression problem, directly relating sequence features to gene expression. Motif Activity Response Analysis (MARA) (Bussemaker et al., 2001; The FANTOM Consortium and Riken Omics Science Center, 2009; Balwierz et al., 2014) is a particularly relevant example of this approach. Conceptually, MARA proposes a model of the following form:

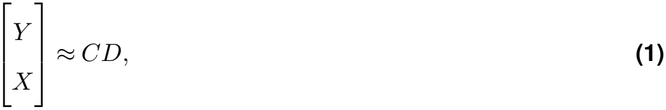

where *Y* and *X* are known gene expression matrices of targets and TFs respectively, with *s* columns corresponding to samples. Thus, rows of *Y* correspond to target genes, whereas rows of *X* correspond to TFs. The reason for making this distinction will become clear later. The known matrix *C* captures sequence features associated with the genes, such as *k*-mer or motif frequencies. The unknown matrix *D* contains the corresponding activities of the sequence features across samples. *D* must be inferred from the expression data. Importantly, the model allows sequence feature activities to vary across samples, such that a distinct linear combination of sequence features contributes to gene expression in each sample. Hence, for each column, corresponding to a particular sample, *D* has an independent set of parameters. The independence between samples provides freedom to account for the potential differences in the presence of TFs across conditions, among other things. Extending this framework into the non-linear realm we find modern Deep Learning approaches such as Enformer, Borzoi, or, more recently, AlphaGenome (Avsec et al., 2021; Linder et al., 2025; Avsec et al., 2026). A relevant limitation of S2E approaches is that, in general, they propose candidate CREs, such as motifs, without identifying the regulatory proteins that recognize them. To attach names to those proteins it is usually required to map the inferred motifs to a database of TFBMs.

Alternatively, the focus can be directed exclusively on the trans-regulatory side by exploiting variation in TF abundance across samples. In this case, cis-regulatory information per gene is disregarded. This is the central premise behind GRN inference: rather than asking how sequence features encode expression, it asks whether variation in TF abundance across samples can explain the variation in target gene expression. This approach typically assumes that TF transcript levels serve as a reliable proxy for the corresponding biologically active protein. Within a linear framework, the canonical regression-based GRN inference model is:

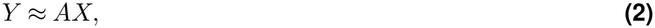

where *A* is an unknown adjacency matrix relating TFs to target genes. Often, this is an under-determined system since there are more TFs than samples. In order to estimate *A*, different regularization approaches are used, such as Least Angle Regression (Haury et al., 2012), Least Absolute Shrinkage and Selection Operator (LASSO) (Omranian et al., 2016) or Network Component Analysis (Liao et al., 2003). When temporal information is available, equation 2 can be formulated as a difference equation, with identifiability addressed through eigen-genes (Alter et al., 2000; Wang et al., 2012; Bansal et al., 2006) or Dynamic Mode Decomposition (Hasnain et al., 2023).

It is common to replace the the linear mapping from TFs to target genes with a non-linear one. Notable examples include random forests, as in GENIE3 (Huynh-Thu et al., 2010), and gradient boosting, as in GRNBoost2 (Moerman et al., 2019), which underpin popular GRN inference pipelines such as SCENIC (Aibar et al., 2017). Regardless of whether the mapping is linear or non-linear, regression-based GRN inference requires fitting a separate model for each target gene. Consequently, applying GENIE3 requites tens of thousands of separate random-forest models. This arises because, within a sample, the abundance of a given TF is shared across all of its potential target genes, whereas the expression of each target gene is distinct. The difference in gene expression within a sample is due to differences on the cis-regulatory elements across genes, which are not taken into account. Hence, regression based GRN inference requires requires gene-specific parameterizations in order to capture the heterogeneous cis-regulatory contexts encoded in their DNA sequences. This is analogous to the S2E situation, where a different model per sample is required due to ignoring trans-elements.

While gene-specific parameterizations provides substantial flexibility, it also obscures the sequence-level structure shared across genes. Moreover, co-expression alone does not imply that the regions regulating a target gene contain a binding site for the corresponding TF. Consequently, current approaches often rely on known, experimentally derived TFBMs to curate inferred regulatory relationships. For example, SCENIC characterizes regulons (sets of genes assumed to be regulated by a particular TF) as filtered subsets of co-expression modules, pruned of spurious targets through motif enrichment analysis (Aibar et al., 2017).

In this work, we propose a novel computational approach that explicitly characterizes the interaction between cis- and trans-regulatory elements; merging GRN and S2E in a single model. We focus on a restricted subset of cis–trans interactions, namely those between TFs and local promoter regions near the transcription start site (TSS). Specifically, we consider a bilinear model of the form:

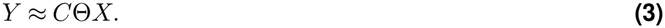

Here, the central unknown is the matrix Θ, which represents the interactions between TFs and genomic sequence features. Our primary objective is to recover Θ as a data-driven approximation of the underlying DNA-binding logic.

Our proposed formulation bridges GRN inference and S2E by explicitly factorizing regulatory effects into a shared interaction matrix Θ between cis-features and TFs. This factorization enables parameter sharing across both genes and samples. In contrast to approaches that fit separate models for individual genes or samples, the regulatory relationships are represented through a common interaction structure. As we show below, the resulting objective is strictly convex and hence has a unique global minimimum, providing a well-defined and interpretable parameterization of the regulatory code. While the present work focuses on a linear formulation, our framework also provides a foundation for future extensions into non-linear regimes, including genomic deep learning.

In the sections that follow, we first demonstrate how a model of the form given in Equation 3, formulated as a bilinear sparse regression, enables the *de novo* inference of TF-specific binding motifs directly from RNA-seq expression data, without requiring further prior experimental knowledge. Second, we show that BRIDGE (**B**ilinear **R**egulatory **I**nference from **D**NA and **G**ene **E**xpression) can predict the expression of unseen promoters in unseen samples, providing a principled and biologically meaningful framework for hyperparameter selection. Finally, we demonstrate that BRIDGE consistently recovers biologically coherent motifs associated with specific TFs across different datasets, many of which closely resemble experimentally validated TFBMs for those TFs.

## Results

### *De novo* discovery of TF-specific binding motifs by integrating transcriptome with promoter *k*-mer frequency vectors

To derive TF–DNA interactions from RNA-seq data and promoter sequence, we propose a regression approach in which the expression of target genes is predicted from TF expression. The main distinction from previous methods is that we force the effect of each TF on a target gene to vary as function of the cis-regulatory elements present in its promoter region. In this way, our model explicitly accounts for the context-dependent nature of gene regulation, in which the regulatory influence of a TF is modulated by the specific cis-regulatory architecture of its target gene. Specifically, we represent the cis-regulatory sequence using 6-mer frequency vectors computed from a local promoter region spanning 500 bp upstream and 500 bp downstream of the TSS. Gene expression is represented using normalized measures such as Transcripts Per Million (TPM), Counts Per Ten Thousand (CPTT), or, when control samples are available, log_2_ fold change with respect of control (log_2_ FC). Other normalization strategies that allow comparisons across genes and cells or conditions may be used. Within the GRN inference framework, this regression problem can be formulated, including intercept terms, as:

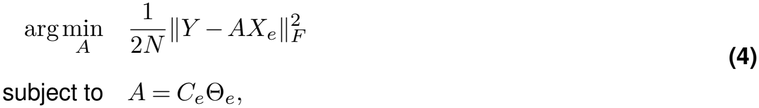

where *N* denotes the total number of observations (target genes × samples). The constraint *A*_*e*_ = *C*_*e*_Θ_*e*_ replaces the unconstrained gene-specific regulatory coefficients of conventional GRN inference with a sequence-dependent parameterization. Here, the subscript *e* indicates that the corresponding matrix has been extended to incorporate intercept terms, such that

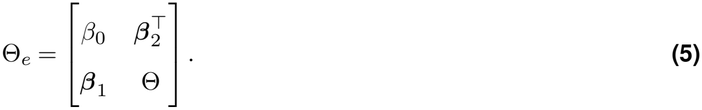

The specific role of the intercept terms is described in the Methods.

A complementary formulation within the S2E framework can be reached following an analogous argument. We aim to predict gene expression based on the presence of cis-regulatory elements, but we constrain the motif activation in a particular sample to be a linear function of TF concentration in that sample. For simplicity, we restrict the dependent variable to be exclusively target genes, alternatively, care must be taken when target genes overlap with transcription factors to prevent the model from learning a trivial identity mapping (i.e. a gene predicting its own expression):

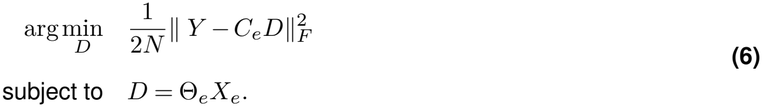

The structure of Θ, however, should reflect the fundamental specificity of gene regulation: each TF interacts only with a limited subset of motifs and therefore we expect Θ to be highly sparse. Hence, our setting can be formulated as the following convex optimization problem (see Methods):

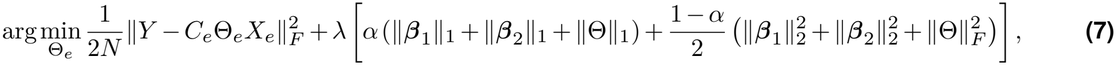

where the global intercept *β*_0_ is not penalized. We propose a FISTA (Beck and Teboulle, 2009) framework and GPU acceleration to efficiently solve the optimization problem outlined in Equation 7, allowing for thorough hyperparameter tuning in little time. Here, we use the classical *L*_1_ penalty because all putative TFs are included in the model. When a curated set of relevant TFs is available, however, it may be advantageous to replace the LASSO component with an “exclusive LASSO” penalty (Campbell and Allen, 2017). The “exclusive LASSO” formulation assumes that the selected TFs are relevant and constrains each TF to interact with only a small subset of sequence motifs. Specifically, the term ∥Θ∥ in Equation 7 can be replaced by 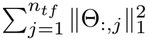, being Θ_:,*j*_ the *j*^th^ column of Θ. This modification can be easily implemented (Lin et al., 2024). A complete description of the optimization algorithm is provided in the Methods.

In summary, given a collection of RNA-seq samples, a list of TFs and a promoter representation, such as matrix of *k*-mer counts, the optimal matrix Θ yields a directly interpretable relationship between TFs and *k*-mers. Aligning the *k*-mers per TF yields a list of pairs of position probability matrices (PPMs) and TFs, which are our putative TF-specific motifs. An overview of the BRIDGE framework is depicted in Figure 1.

**Figure 1.**
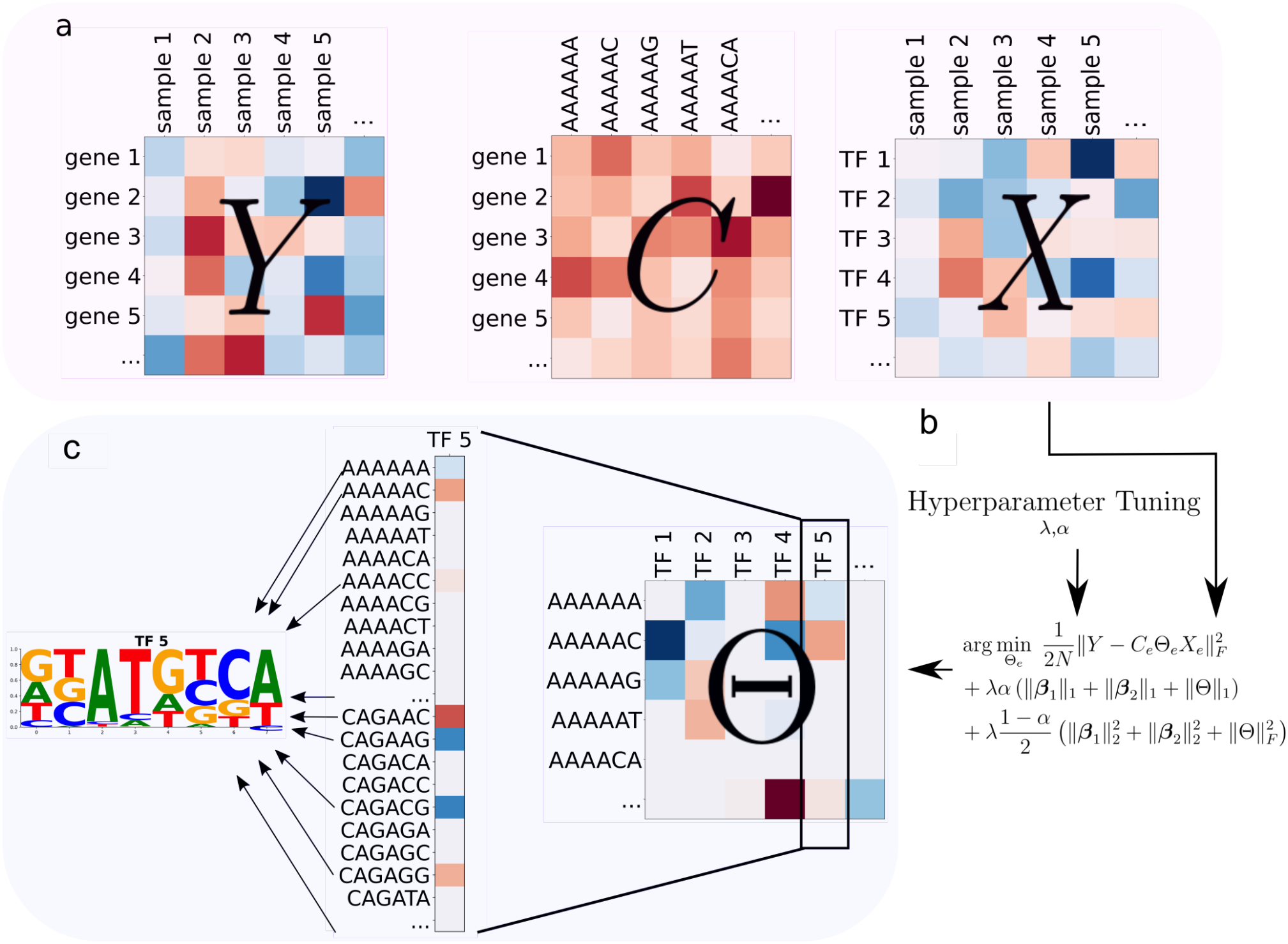
A Bilinear Sparse Regression framework for de novo TFBM discovery. **a)** The BRIDGE framework integrates a normalized gene expression count (e.g.log(1 + TPM)) matrix *Y*, promoter *k*-mer frequency matrix *C* and TF expression count matrix *X*. **b)** Θ_*e*_ minimizes the error in the approximation of the gene expression of target genes using information from the genome and expression of TFs, while respecting the elastic-net induced sparsity. **c)** The resulting weight matrix Θ maps specific *k*-mers to the regulatory influence of each TF; individual columns of Θ represent *k*-mer sets that, when aligned, yield putative TFBMs.

### Modeling TF-DNA interactions improves predictive performance in unseen genes and unseen conditions

Traditionally, predictive performance of GRN inference algorithms is evaluated in held-out samples or cells. This approach is, however, problematic. It cannot discern between co-expression and regulation, which is a well-known issue.

Gene expression prediction in held-out genes, even within seen samples is a much harder and meaningful task. By design, BRIDGE predicts unseen target genes based on their promoter and unseen conditions based on the TFs concentrations. Hence, model validation in BRIDGE is based on predictive performance in unseen samples and unseen genes. Within that setting, we evaluated the added value of explicitly modeling interactions between 6-mers and TFs as opposed to relying solely on their independent contributions. Indeed, if the inferred TFBMs are biologically meaningful, explicitly modeling these interactions should yield a measurable improvement in predictive performance in test set. Note that, for the same hyperparameters, the full model will always achieve better predictive performance in the training set. However, here we are concerned with actual generalization performance and therefore we focus exclusively on unseen samples and genes.

To rigorously assess the utility of formally capturing the TF-DNA interaction we fitted two models. The full model includes a global intercept *β*_0_ (constant across genes and samples), main effects for 6-mers *β*_1_ (constant across samples), main effects for TFs *β*_2_ (constant across genes) and interaction terms Θ between TFs and 6-mers. A restricted version of that model includes only the global intercept and the main effects for TFs and 6-mers. This is easily accomplished by forcing Θ = 0 within Θ_*e*_ in Equation 7, following the same optimization procedure. The restricted model assumes that the effect of a 6-mer in expression is the same regardless the cellular state and, analogously, that the effect of the presence of a particular TF in the expression of a gene is independent of the cis-regulatory characteristics of its promoter. Clearly, the restricted model violates fundamental characteristics of gene regulation and thus is highly unrealistic, but it serves as a baseline to study the effect of including cis-trans interactions. For both models hyperparameter tuning and testing is conducted in exactly the same way (see Methods).

We tested the full model and its restricted version on 8 different transcriptome datasets of human and *Arabidopsis thaliana*, spanning bulk (Bulk), single-cell (SC) and pseudo-bulk single-cell RNA-seq (PBulk) data. In all cases, the counts are normalized to log Transcripts per Million (log(1 + TPM)) or its similar singe cell version log Counts per Ten Thousand (log(1 + CPTT)). Furthermore, leveraging the availability of structured experiments for *Arabidopsis thaliana*, where a control per time point and condition is available, we included a different normalization approach based on log_2_ fold change (log_2_ FC) with respect to control. log_2_ FC variability corresponds more clearly to condition-specific changes, for example, ignoring transcriptomic patterns involved in circadian rhythm. An overview of the dataset characteristics, numbers of samples, genes and TFs is provided in Table 1; a more detailed description of the datasets and dataset processing is provided in the Methods.

**Table 1.** Summary of dataset characteristics, structural dimensions, sparsity levels, and optimized hyperparameters. Ath stands for *Arabidopsis thaliana*. “Data Type” is bulk RNA-seq (Bulk), single-cell (SC) or SC aggregatted into pseudo-bulk (PBulk). For each dataset, the number of different conditions, tissues, cells and/or replications is represented as “Samples”. The number of target genes is listed in the “Genes” column, the number of TFs in the “TFs” column. “Sparsity” refers to the % of 0s in Θ_*e*_. “Active TFs” refers to the number of TFs that have at least one active (non-zero) 6-mer associated in the full model. Similarly, “Active 6-mers” refers to the number of active 6-mers that are present in at least one TF. “Act. Int. TFs.” and “Act. int. 6-mers” refers to the non-zero main effects for TFs and 6-mers respectively. The selected hyperparameters for the full model are displayed in the *α* and *λ* columns.

| Dataset | Species | Data Type | Samples | Genes | TFs | Sparsity (%) | Active TFs | Active 6-mers | Act. int. TFs | Act. int. 6-mers | α | λ |
|---|---|---|---|---|---|---|---|---|---|---|---|---|
| PTI (TPM) | Ath | Bulk | 287 | 22,448 | 1,649 | 99.98 | 13 | 678 | 0 | 0 | 0.905 | 0.050 |
| Hormone (TPM) | Ath | Bulk | 276 | 22,837 | 1,671 | 99.91 | 80 | 798 | 0 | 0 | 0.021 | 1.597 |
| Leaf (CPTT) | Ath | SC | 4,707 | 17,041 | 1,188 | 99.96 | 135 | 551 | 0 | 65 | 0.471 | 0.005 |
| PTI (log<sub>2</sub> FC) | Ath | Bulk | 265 | 22,448 | 1,649 | 99.90 | 404 | 839 | 0 | 17 | 0.305 | 0.008 |
| Hormone (log<sub>2</sub> FC) | Ath | Bulk | 225 | 22,837 | 1,671 | 99.92 | 292 | 767 | 0 | 0 | 0.574 | 0.013 |
| SN Hormone (log<sub>2</sub> FC) | Ath | PBulk | 389 | 25,529 | 1,655 | 99.69 | 868 | 1,471 | 0 | 0 | 0.622 | 0.003 |
| SN Hormone (CPTT) | Ath | PBulk | 540 | 25,554 | 1,657 | 99.98 | 39 | 432 | 0 | 0 | 0.166 | 0.023 |
| GTEx (TPM) | Human | Bulk | 17,654 | 42,357 | 1,630 | 99.96 | 99 | 745 | 0 | 0 | 0.475 | 0.042 |
| Tabula Sapiens (CPTT) | Human | PBulk | 21,366 | 44,480 | 1,636 | 99.94 | 159 | 670 | 0 | 3 | 0.879 | 0.001 |
| PBMC (CPTT) | Human | SC | 9,252 | 16,320 | 1,262 | 99.98 | 18 | 322 | 0 | 0 | 0.039 | 0.175 |
| TCGA (TPM) | Human | Bulk | 9,481 | 19,337 | 729 | 99.87 | 154 | 520 | 0 | 0 | 0.379 | 0.383 |

After optimizing hyperparameters through cross-validation (CV) on the training set, model performance was evaluated on a held-out set composed of unseen genes and unseen samples. Note that for any given hyperparameter configuration, the optimization yields a unique solution (see Methods). The predictive performance of the full and restricted model across each dataset is summarized in Figure 2. To provide a clear idea of predictive performance, we break down the variance explained (*R*^2^) metric into three different views: first, the *R*^2^ naturally arising when comparing the ground truth vec(*Y* ) and the prediction vec(*C*_*e*_Θ_*e*_*X*_*e*_) (panel a); second, the average *R*^2^ per gene across samples (panel b); and third, the average *R*^2^ per sample across genes (panel c). The averages are calculated with negative individual *R*^2^ values trimmed down to 0. This breakdown is interesting because, depending on the normalization strategy, the variance of samples may be considerably higher than the variance of genes. In such cases, the overall *R*^2^ may not reflect the transcriptome dynamics being modeled well, but rather somewhat static levels of expression per gene. Several studies show that predicting average levels of expression is considerably easier than predicting gene expression dynamics across different conditions (Alonso Esteve et al., 2026; Peleke et al., 2024; Washburn et al., 2019).

**Figure 2.**
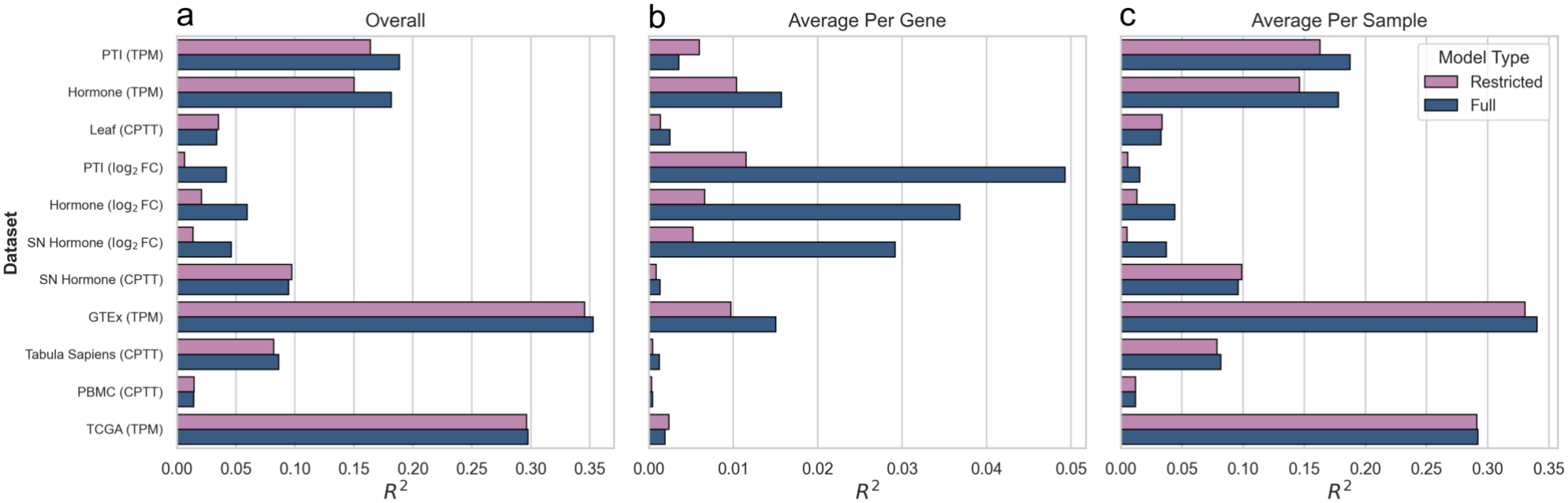
Predictive performance on unseen samples and genes: comparison between the restricted and full model. The variance explained (*R*^2^) by the predictions is evaluated at three levels of granularity. In panel **a)**, “Overall” *R*^2^ represents the total variance explained across all genes and samples in the entire test set. In panel **b)** “Average Per Gene” and panel **c)** “Average Per Sample” represent the *R*^2^ averaged across all samples for each individual gene, and across all genes for each individual sample, respectively. The *R*^2^ per gene relates to how well the model captures the variability of a gene across conditions, while the *R*^2^ per sample relates to how well the differences in expression between genes are captured.

Overall, predictive performance improves with the inclusion of interaction terms in all but the “raw” single-cell datasets and the pseudo-bulk hormone dataset with CPTT normalization, with a decrease in *R*^2^ of 3.5%, 1% and 2.8% for Leaf (CPTT), PBMC (CPTT) and SN Hormone (CPTT) data respectively. In both bulk *Arabidopsis thaliana* datasets, Hormone (TPM) and PTI (TPM), there is an increase in predictive performance, of 15.5% and 21% respectively. This increase is even clearer when using log_2_ FC normalization, although with a large decrease in *R*^2^ to TPM. Similarly, while the absolute *R*^2^ decreases with log_2_ FC normalization in the single-cell pseudo-bulk hormone dataset, the inclusion of interaction terms in this case has a positive effect in predictive performance. This information is available in Table S1.

As seen in Figure 2b, the *R*^2^ in TPM- and CPTT-normalized data is largely due to relatively accurate prediction of the variation across genes, but a very poor prediction of the dynamics per gene. For the PTI (TPM), Hormone (TPM), GTEx (TPM) and TCGA (TPM) datasets (Figures S1, S2, S3, and S4) we find that the variability of expression within a sample tends to be higher than the variability of a gene across samples. This explains the similarity of the overall (panel a) and average per sample (panel c) predictive performance and suggests that the “scale” of expression per gene is captured better than the condition-specific variability.

In this context, results obtained after log_2_ FC normalization are particularly relevant. In this case, the standard deviation distributions of genes and samples are much closer (Figures S6, S5, S7), which is reflected in the similarity of the *R*^2^ across all three views (panels a, b and c in Figure 2). Furthermore, not only is the predictive performance per gene superior to that on TPM- and CPTT-normalized data in absolute terms, but the increase due to inclusion of interactions is greater as well.

Inspection of the optimal Θ_*e*_ for the BRIDGE model reveals a clear preference for very sparse solutions across all datasets, as expected. As shown in Table 1, sparsity is above 99.5% in all cases. Furthermore, intercept effects become 0 (column “Act. Int. TFs” and “Act. Int. 6-mers” in Table 1) across the different datasets with minor exceptions, suggesting that indeed, the interaction between 6-mers and TFs is overall more relevant for prediction than their main effect.

The distributions of 6-mers per TF, however, display long tails (Figure S8). Ideally, active TFs would bind a small, coherent subset of 6-mers. However, we observed a handful of outlier TFs predicted to interact with hundreds of sequence features. For example, in the PTI (TPM) dataset, EBP (AT3G16770) is connected to roughly 700 6-mers. While this broad profile might initially appear to lack binding specificity, the estimated main effect (constant term) for EBP is zero. The fact that the model preferred a distributed linear combination of sequence features over a simple non-zero baseline suggests that this broad activation reflects genuine, nuanced regulation rather than non-specific background noise and that EBP may act as a systemic regulator in defense- and stress-related responses.

Conversely, the distribution of 6-mer occurrences across TFs displays a shorter tail (Figure S9). The outliers in this case are driven by prevalent AAAAAA-like motifs that associate with more than 100 TFs in some cases. This might be related to chromatin accessibility (Cuesta-López et al., 2011).

### Learned TF-DNA interactions reconstruct experimentally validated motifs

With the optimal sparse Θ at hand, we can proceed to inspect what 6-mers a particular TF is predicted to bind. As outlined in Figure 1d, each column in Θ gives some non-zero weights to a subset of 6-mers. We use these selected 6-mers to construct candidate TFBMs for each active TF. Using an importance score (see Methods) associated to the Θ_*i,j*_ parameter for the *i*^th^ 6-mer and *j*^th^ TF, we construct candidate TFBMs by optimally aligning each active 6-mer to the most important one, which we call the anchor 6-mer (see Methods). The resulting PPM for each TF is a weighted sum of each 6-mer with optimal position relative to the anchor 6-mer.

To see if the obtained candidate TFBMs are indeed correct, we now consider their similarity to the JASPAR (Ovek Baydar et al., 2026) counterparts. For each TF we check if our proposed candidate TFBM is similar to the experimentally derived one, obtained from the JASPAR database. Figure 3 outlines the results of an (TOMTOM) analysis of our BRIDGE-derived TFBMs against a (JASPAR) database. Figure 3a depicts the QQ-plot of −log_10_ p-values under the null hypothesis of no relevant information gathered vs the actual raw TOMTOM −log_10_ p-values for specific TFBMs. Under the null, points are expected to be along the diagonal. Deviation above the diagonal indicates an excess of low p-values relative to expectation, reflecting significant enrichment of motif matches. Note that the comparison against the expected uniform distribution is done independently for each dataset.

**Figure 3.**
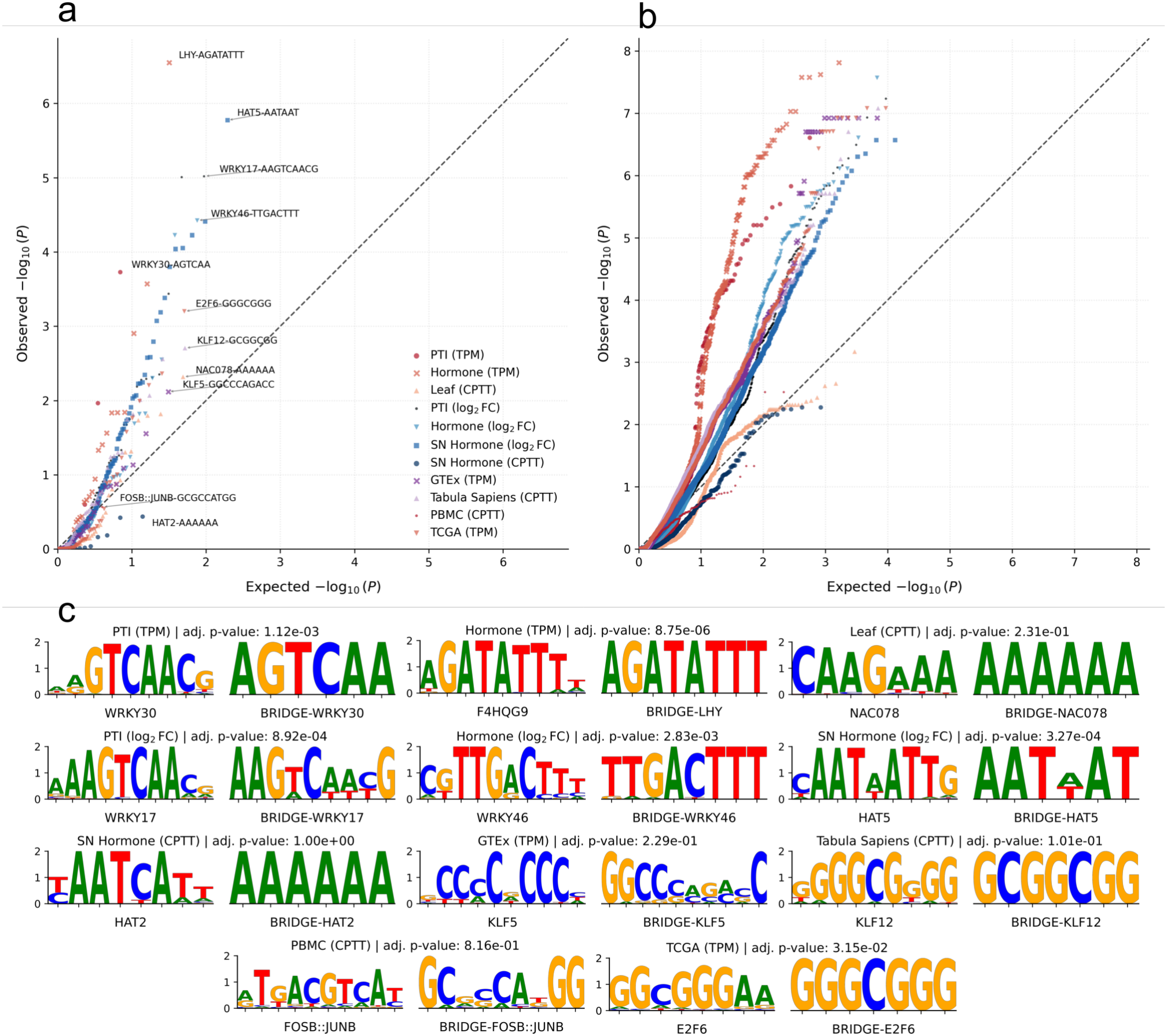
Comparison of inferred TFBMs with JASPAR experimentally validated counterparts. **a)** Expected, uniformly distributed, −log_10_ p-values under the null hypothesis of absence of biological signal in the horizontal axis vs. actual raw TOMTOM −log_10_ p-values for specific TFBMs. Each inferred TFBM is compared exclusively with its experimentally derived one, no cross-comparisons are performed. Strong deviation from the diagonal line indicates significant recovery of TFBMs. The best scoring TFs per dataset are listed, with their inferred consensus motif. **b)** Inferred TFBMs compared with all available experimentally derived counterparts within their structural family. Since DNA Binding Domains (DBD) are heavily conserved, these similarities still recover the biological relevance of the inferred TFBMs but are more forgiving to small deviations and allow evaluation for TFs not present in JASPAR. **c)** The top scoring inferred TFBMs (on the right) and their JASPAR counterpart (on the left). These TFs are named in panel a. The Bonferroni corrected (over the TFs considered in a particular dataset) raw TOMTOM p-values are provided.

Evaluating predicted motifs strictly through one-to-one matching with specific reference TFBMs presents a significant challenge: subtle positional deviations or variations in motif width can yield severe matching penalties for otherwise coherent predictions. Furthermore, experimental reference motifs in databases like JASPAR may not be linked to specific TFs, even when extensive binding profiles exist for structural paralogs. While Figure 3a shows one-to-one results, Figure 3b displays a one-to-many relationship. For each TF considered, panel b shows the raw TOMTOM −log_10_ p-value against each element within that TF’s family available in JASPAR. Figure 3b leverages the high sequence and structural conservation of DNA Binding Domains (DBDs) within TF families. By evaluating motif similarity across all members of a structural family, we account for positional noise and reference incompleteness without unfairly penalizing biologically valid inferences. Note that, the datasets for which the recovery is very poor in Figure 3a, display a clear pattern along the diagonal on Figure 3b, for instance Leaf (CPTT), SN Hormone (CPTT) or PBMC (CPTT).

It is worth noticing that this is fundamentally different from the typical enrichment strategy where clustering is followed by a motif discovery algorithm such as STREME or MEME (Bailey, 2021; Bailey et al., 1994). Enrichment analyses are based on the over-representation of some motifs within a particular group of, for instance, differentially expressed genes in a particular condition. However, BRIDGE is, fundamentally, a predictive model. BRIDGE finds TFBMs by minimizing the reconstruction loss of the transcriptome landscape conditional on promoter features and TF concentrations. Because BRIDGE does not require clustering as a precondition, the inferred motifs are TF-specific, as can be seen in Figure 3a and c.

Overall, Figure 3a and b, show that BRIDGE recovers relevant biological signal, as can be appreciated in some of the instances shown in panel c. For example, the BRIDGE-inferred LHY (UniProt ID: F4HQG9) TFBM shows striking similarity to the actual JASPAR motif, which is interesting given that the inferred motif is derived solely from RNA-seq and DNA sequence data. The best reconstructed motifs are exclusively found for the *Arabidopsis thaliana* datasets. In particular, the WRKY TF family, known to be highly relevant for stress responses in *Arabidopsis thaliana* (Bjornson et al., 2021), seems to be the easiest to reconstruct. The reconstruction for the human datasets is worse and the reconstruction for the CPTT normalized single-cell data, in both species, appears difficult. From Figure 3b, it is very clear that for SN Hormone (CPTT), PBMC (CPTT) and Leaf (CPTT), no relevant signal is recovered.

The capabilities outlined above enable the generation of putative TFBMs directly from transcriptomic and sequence data, opening the door to characterizing previously unannotated TFs by repurposing existing RNA-seq datasets. However, given the large number of potential candidates this yields, a clear prioritization strategy is essential for experimental follow-up. TFs with high assigned importance score should be, ideally, the most biologically plausible ones. Figure 4 illustrates the relationship between the importance score assigned to the anchor 6-mer for each TF and its TOMTOM similarity −log_10_ pvalue. Overall, the relationship is again strongest for the Arabidopsis datasets and particularly weak for the human datasets. When looking instead at the best match per family (see Figure S11), we find a significant (*p <* 0.001) Spearman correlation of 0.1.

**Figure 4.**
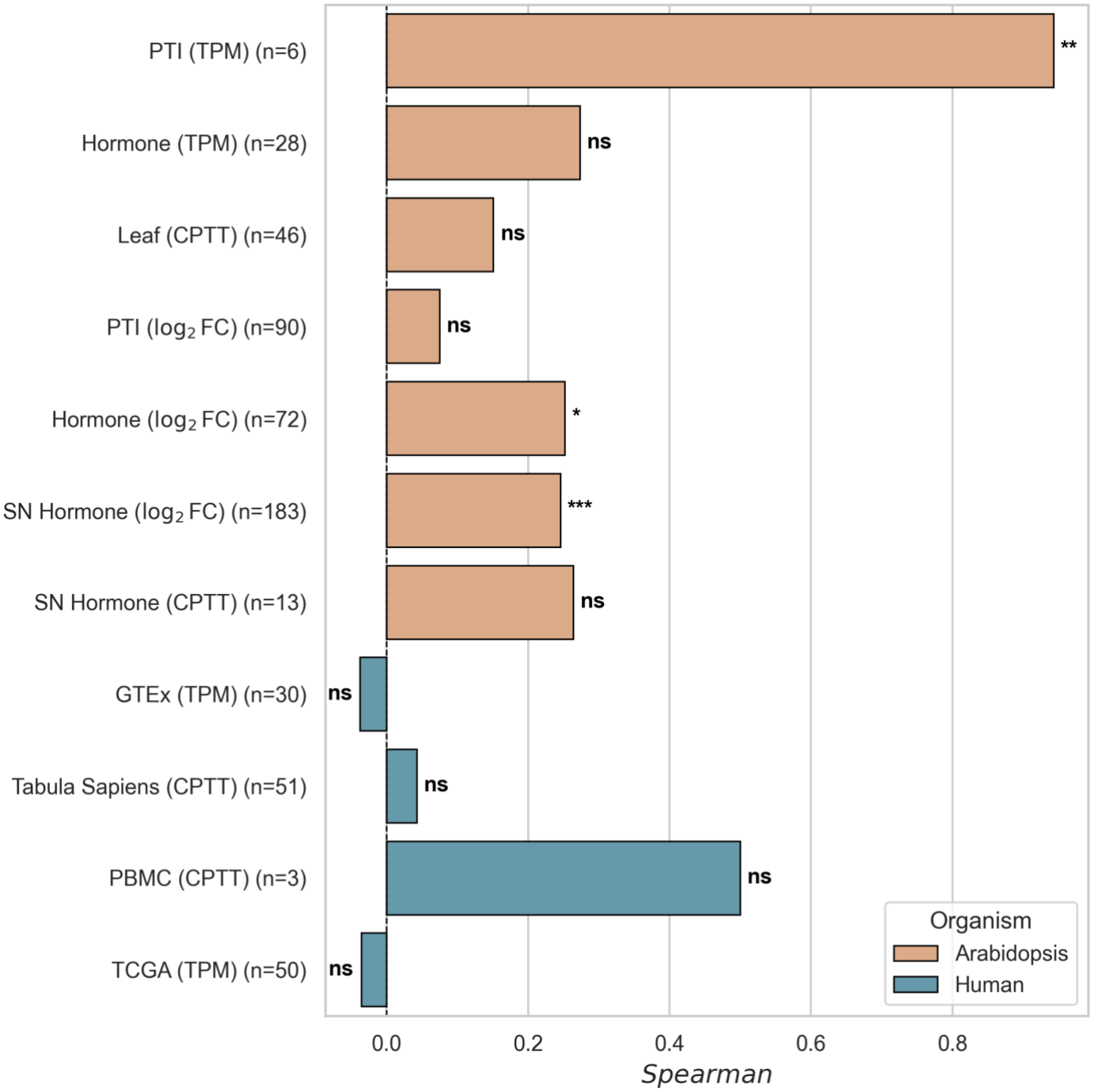
Spearman correlation between the importance scores of the anchor 6-mer and the TOMTOM raw −log_10_ p-values per TF. For each dataset, the relationship between inferred TFBM importance and its similarity to the experimentally derived one is evaluated. The number of TFs considered is displayed besides the dataset name. Note that there is a reduced number of TF that pass the filter IC filter to be considered, and here that number is further reduced to the TFs that are also present in JASPAR. * *p <* 0.05, ** *p <* 0.01, *** *p <* 0.001, ns = not significant.

The most important inferred TFBMs per dataset for the subset of TF that are not characterized in JASPAR can be seen in Figure S10, offering high-priority candidates for targeted experimental validation. The complete set of BRIDGE-inferred TFBMs per dataset is provided in .meme format in the supplementary material.

## Discussion

In this work, we introduced BRIDGE, a framework that unifies GRN inference and S2E modeling into a single, convex optimization problem. By explicitly factorizing regulatory interactions into a shared matrix Θ mapping *k*-mers (or any other promoter representation) to TF abundances, BRIDGE parameterizes regulatory interactions and guarantees a unique global solution. Beyond predictive accuracy, this formulation yields direct biological interpretability: individual regulatory edges in the reconstructed GRN (*C*Θ) can be dissected to reveal their sequence-level motif basis. The mathematical simplicity and computational efficiency of the proposed method allows it to serve as a baseline model for gene expression prediction.

We have shown that including interactions between 6-mers and TFs tends to improve predictive performance on unseen genes and unseen samples. This is expected, as gene regulation is a highly dynamic process dominated by the interaction of cis- and trans-regulatory elements. However, the predictive performance of BRIDGE is not yet sufficient for applications such as promoter sequence engineering (Sadravi et al., 2025). In the best cases, average *R*^2^ reaches 0.05. This is not surprising: while the simplicity of the model allowed to isolate the effect of the cis-trans interaction, it builds on several limiting assumptions. These limitations include ignoring cooperative binding of TFs, ignoring positional information of the regulatory elements, and linearity (e.g. doubling the TF expression level doubles its effect). Moreover, using a promoter region between 500bps upstream and 500bps downstream TSS may be suboptimal for human datasets. While *Arabidopsis thaliana* has a very compact genome, human gene regulation is more complex and relies more heavily on distal regulators, which are not captured here. Furthermore, human datasets comprise different genomic backgrounds mapped to the same reference genome, which adds an additional source of noise relative to the *Arabidopsis thaliana* experiments considered. Some of these limitations could be addressed, at the expense of simplicity and interpretability, through smart applications of genomic deep learning or other approaches such as tree-based methods. However, this is not a simple task, as more powerful models do not necessarily provide better predictive performance when data is scarce (Alonso Esteve et al., 2026).

Throughout this work, we have assumed a static genomic sequence and constant chromatin state. Nevertheless, dynamic epigenetic mechanisms exert profound control over transcription factor occupancy and gene regulation (Klemm et al., 2019; Kouzarides, 2007). Interestingly, BRIDGE could be extended to deal with varying chromatin states simultaneously by allowing for 3-dimensional arrays for *C, Y* and *X* while preserving the very intuitive 2D structure in Θ. Interesting datasets that would allow this, for instance, couple ATAC-seq with RNA-seq single-cell sequencing (Ma et al., 2020); however, as we have seen, “raw” single-cell RNA-seq remains challenging to model.

A central assumption of our current framework is that TF transcript abundance serves as a direct proxy for biologically active protein concentration. If transcript levels diverge significantly from active protein states, the capacity of the model to accurately infer true regulatory influence is reduced. The fact that we can recover some interesting TF-specific binding motifs from just RNA-seq data with BRIDGE, albeit modestly, suggest that this assumption is not totally misplaced.

Biologically, we expect each TF to bind a small, coherent subset of 6-mers, so the relevant sparsity is structured (few motifs per TF, and few TFs overall) instead of arbitrary. While we have shown that element-wise LASSO is an effective regularization strategy, it treats all entries of Θ independently and cannot encode this structure. Consistent with this, the distribution of 6-mers and TFs within Θ shows room for improvement: several outlier TFs bind many motifs, in some cases above 500. Combining “exclusive LASSO” (Campbell and Allen, 2017) with “group LASSO” (Yuan and Lin, 2006) could enforce a trade-off (Liu et al., 2019) in which entire TFs are excluded from the model altogether, while among the TFs that are retained, competition is imposed between their associated motifs so that only a sparse subset of coherent 6-mers is selected per TF.

Finally, BRIDGE assumes *k*-mers and TFs to be all independent entities; however, *k*-mers have varying degrees of similarity and TFs have conserved DNA-binding domains within their families. A possible extension could account for similarities between *k*-mers using, for example, a Laplacian penalty (Li et al., 2010), which can be very easily integrated, as it is convex and differentiable function. TF families or structural similarities could be integrated in a similar fashion.

In conclusion, BRIDGE demonstrates that even within the constraints of a simple linear formulation, explicitly modeling cis–trans interactions exposes rich biological signals that are often lost in pure co-expression or sequence-only approaches. The transparency of the method provides a clear diagnostic lens to evaluate data quality, normalization choices, and modeling assumptions. As regulatory genomics moves toward integrating single-cell dynamics, spatial accessibility, and deep learning, BRIDGE offers a principled conceptual foundation for future multi-omics and non-linear regulatory models to be built on.

## Methods

### Bilinear sparse regression

Let 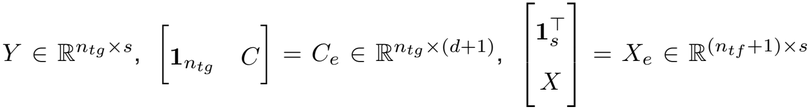 be known, intercept extended, data matrices of target genes expression, promoter representations such as 6-mer counts and TF expression respectively, with *n*_*tg*_ the number of target genes, *n*_*tf*_ the number of TFs, *d* the number of dimensions of the promoter representation and *s* the number of samples, e.g. experimental conditions or cell lines. The matrix 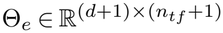 is an extended unknown matrix of TF-genome interactions. The extra dimension in the number of genomic features and TFs accounts for main effects and intercept terms:

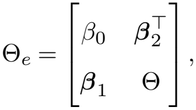

where 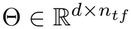 is the TF-DNA interaction matrix, *β*_0_ is a scalar acting as global intercept, ***β***_1_ ∈ ℝ^*d*×1^ and 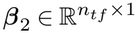 are the main effects of *k*-mers and TF respectively.

The optimization problem of Equation 7 is solved through an application of FISTA, implemented in JAX (Bradbury et al., 2018) for GPU parallelization of linear algebra operations. FISTA is designed to work for an objective function of the form min_*x*_{*F* (*x*) ≡ *f* (*x*) + *g*(*x*)}, *x* ∈ ℝ^*n*^ where the function *f* is a convex, continuously differentiable function and with constant *L*(*f* ) such that

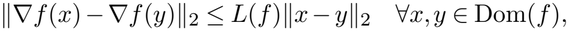

and function *g* is continuous and convex, but not necessarily differentiable. It is easy to see how the bilinear sparse regression setting fits those requirements by noting that Equation 7 is just Ordinary Least Squares (OLS) with an elastic-net penalty. In what follows we show this explicitly.

Since “flattening” any matrix and applying the *L*_2_ norm is the same as the Frobenius norm on that matrix, we have the following equivalent objective function in vector form using the Kronecker product:

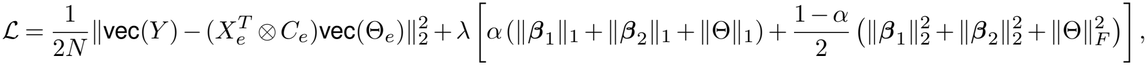

which is the canonical elastic-net formulation (Zou and Hastie, 2005). Importantly, although mathematically equivalent, the OLS formulation of this problem is computationally impractical due to the size of 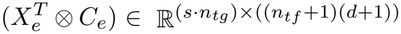 even for modest number of dimensions, TFs and/or samples.

Looking at the gradient of the smooth part (ignoring the *L*_1_ penalty) we find:

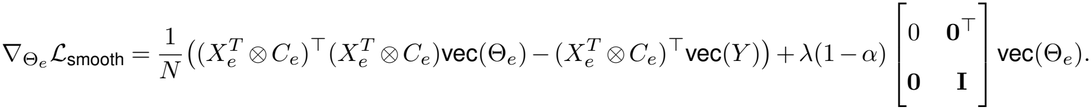

We want to show that this function has a Lipschitz continuous gradient. We have:

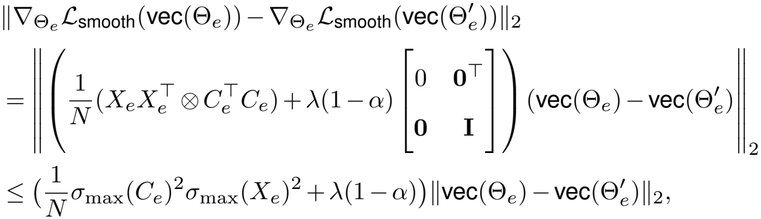

where the last step follows from Weyl’s inequality, 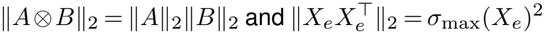, for *σ*_max_(*C*_*e*_) and *σ*_max_(*X*_*e*_) being the maximum singular value of *C*_*e*_ and *X*_*e*_ respectively. Therefore ℒ_smooth_ fulfills the requirement of having a Lipschitz continuous gradient.

Next, we look for convexity. For the Hessian of the smooth part we have:

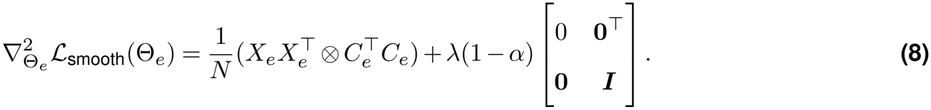

Since the eigenvalues of a Kronecker product are the product of the eigenvalues of each matrix and the Gram matrices are always positive semi-definite, the first term is positive semi-definite. To show strict convexity and hence uniqueness of the solution we need to show positive definiteness of the Hessian. For *α <* 1 and *λ >* 0 (which holds for every model fitted in this work) we have that for all vectors 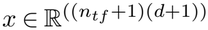:

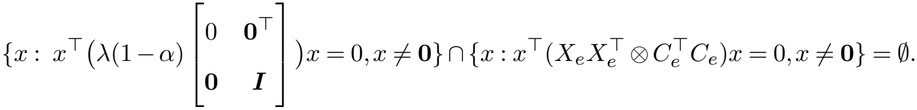

Clearly, the first set is constructed by vectors with non-zero element on the first entry and zeros otherwise, i.e. {*ce*_1_ : *c* ≠ 0}, where *e*_1_ is the first standard basis vector. Now, those vectors will never be in the nullspace of 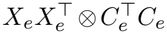 since the first entry of that matrix is *s* · *n*_*tg*_ = *N* . Hence, the smooth part is a strictly convex function.

In summary, ℒ_smooth_ is a strictly convex function with a Lipschitz continuous gradient and hence a valid candidate for *f* in FISTA.

For the non-smooth part we have

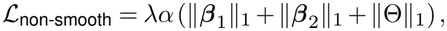

which is convex and non-smooth. This is a valid candidate for *g*. Note that the sum of a strictly convex function and a convex function yields a strictly convex function.

Since the Hessian of ℒ_smooth_ is constant and positive definite (so it is coercive) and ℒ_non-smooth_ ≥ 0, Equation 7 has a minimizer, which is unique by strict convexity. Having established that ℒ = ℒ_smooth_ + ℒ_non-smooth_ satisfies the conditions required by FISTA (Beck and Teboulle, 2009), we leverage it to solve Equation 7. FISTA guarantees *F* (Θ_*k*_) − *F*^*⋆*^ = *O*(1*/k*^2^), where *k* is the number of iterations and *F*^*⋆*^ the minimum of the loss.

The implementation is described in Algorithm 1.

#### Algorithm 1

FISTA on Bilinear Sparse Regression

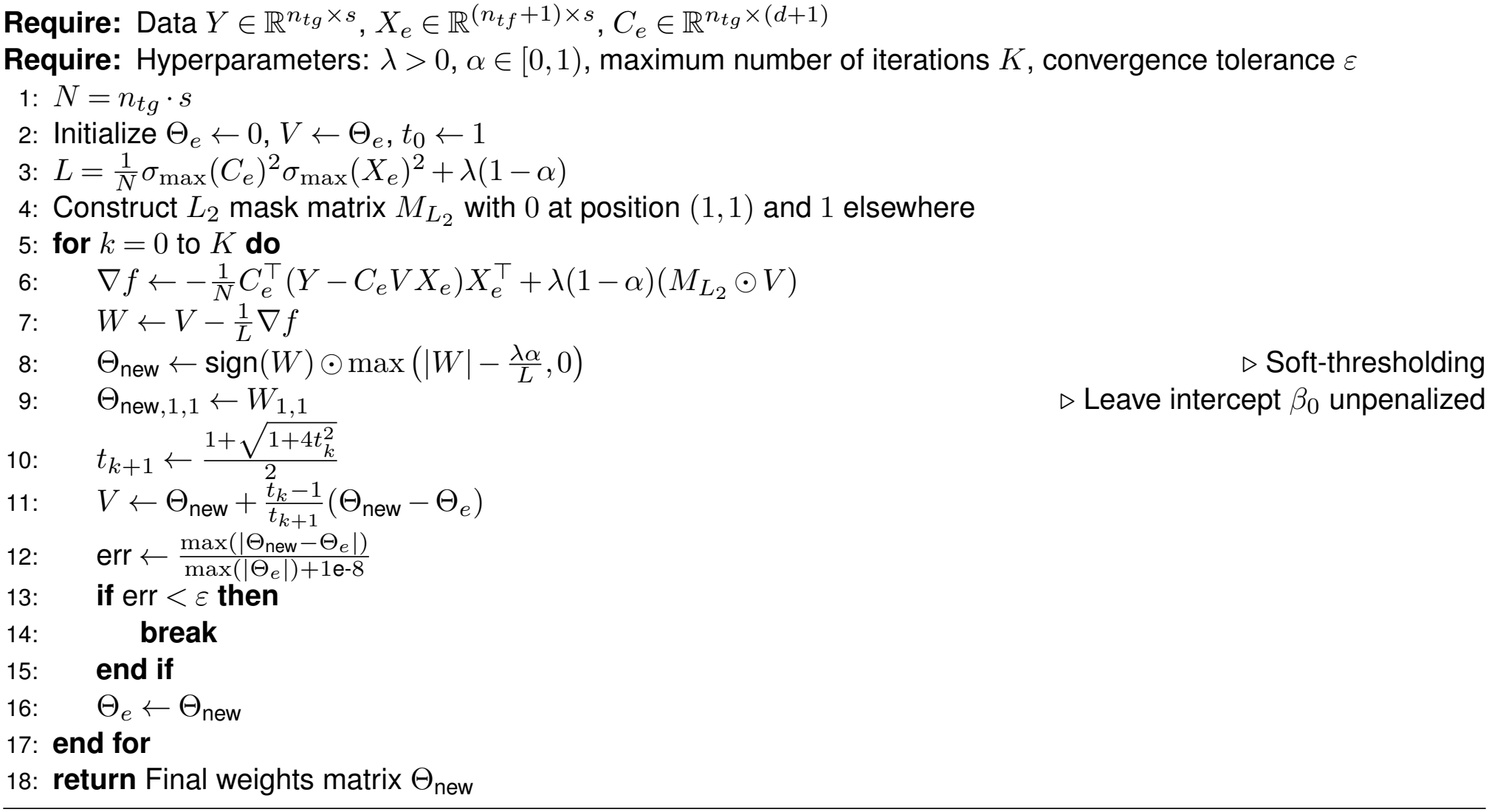

The restricted model without interactions, i.e. with parameter 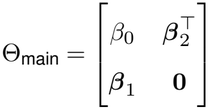, was optimized using algorithm 1 but enforcing all elements in Θ to be 0.

### DNA data processing

Genome sequences and annotation for *Arabidopsis thaliana* were obtained from the TAIR website (Rhee et al., 2003). We used the TAIR10 genome sequence and annotation, with the exception of the single-cell hormone data, where the Araport11 annotation was used (Lamesch et al., 2012). For human, GRCh38 genome sequence data was obtained from Ensembl and annotation from GENCODE (Kinsella et al., 2011). Gene annotations were filtered to retain only HAVANA-curated gene models. All genes are considered, but only the intersection with the genes present in the RNA-seq data will be used. Formatting was applied to ensure that the .fasta and .gff files of both species were usable for downstream pipelines.

To generate the matrix *C* of genes by *k*-mers we fixed *k* = 6 and 500 bp up- and downstream of the TSS. For each gene, the number of times a particular 6-mer appears in that 1000bp window is recorded in a vector of 6-mer counts. We assume reverse complement equivalence, and therefore appearance counts of the forward and reverse complement of a particular 6-mer are summed. Once the process is done for all genes, we end up with as many vectors of 6-mer counts as genes in the annotation file and the matrix *C* with 2080 columns is ready. 2080 corresponds to the number of canonical 6-mers after accounting for reverse complement equivalence. In the 4^6^ = 4096 possible 6-mers, palindromic 6-mers are counted once rather than twice. To get the total number of unique 6-mers (up to reverse complement) we need to add them again before dividing by 2, giving (4^6^ + 4^3^)*/*2 = 2080 (three bases are free in 6-mer palindromic sequences) distinct 6-mers.

### RNA-seq data processing

In the table 2 the links to download each dataset are provided, with their corresponding format. Note that a guide for downloading the data and reproducing the results is provided in the code repository.

**Table 2.** Overview of Datasets, Species, Normalization, and Sources.

| **Dataset** | **Species** | **Normalization** | **Link** |
|---|---|---|---|
| PBMC (Zheng et al., 2017) | Human | log(CPTT + 1) | 10x Genomics |
| Leaf (Kim et al., 2021) | *Arabidopsis thaliana* | log(CPTT + 1) | Plant SC browser |
| SC Tabula Sapiens (The Tabula Sapiens Consortium, 2022) | Human | log(CPTT + 1) | CellxGene |
| Hormone (Aerts et al., 2024) | *Arabidopsis thaliana* | log(TPM + 1) & log<sub>2</sub> FC | Article |
| PTI (Bjornson et al., 2021) | *Arabidopsis thaliana* | log(TPM + 1) & log<sub>2</sub> FC | Article |
| TCGA (Weinstein et al., 2013) | Human | log<sub>2</sub>(TPM + 0.01) | Xenabrowser |
| GTEx (Lonsdale et al., 2013) | Human | log(TPM + 1) | GTEx Portal |
| SN Hormone (Liu et al., 2026) | *Arabidopsis thaliana* | log(CPTT + 1) & log<sub>2</sub> FC | Article |

The TCGA (The Cancer Genome Atlas) (Weinstein et al., 2013) dataset already comes normalized in log_2_(TPM + 0.01) and we used it as is. The GTEx (The Genotype-Tissue Expression) (Lonsdale et al., 2013) dataset comes in TPM and we simply transformed it into log(TPM + 1). For the Hormone and PTI (Aerts et al., 2024; Bjornson et al., 2021) datasets, raw mapped gene counts were converted to log(TPM + 1) values using the following procedure: For a given gene *i* with total exon length *L*_*i*_ (in kilobases) and raw count *C*_*i,j*_ in sample *j*, Reads Per Kilobase (RPK) were first calculated as:

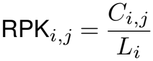

These values were subsequently normalized by the total depth scaling factor of sample *j* to yield TPM:

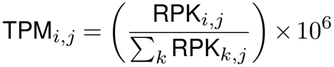

where the denominator represents the sum of RPK values across all genes *k* in sample *j*. For log_2_ FC normalization, first we applied DEseq2 (Love et al., 2014) median of ratios normalization so samples can be compared. For each condition and time point, control replicates were averaged to establish a baseline expression level 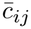 for gene *i* and sample *j*. Relative gene expression changes for each treatment sample *t*_*ij*_ were quantified as the log_2_ fold change relative to its matched control:

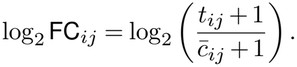

Values were clipped to [−4, 4]. In both cases, genes with zero standard deviation in expression across all samples were removed prior to downstream analysis.

For the SN Hormone dataset (Liu et al., 2026), pseudo-bulk samples were generated by summing up the cell samples by hormone × time point × cell type (as outlined in their manuscript). For SC Tabula Sapiens dataset (The Tabula Sapiens Consortium, 2022) we followed the procedure from Quake et al. (2025). For each tissue and cell-type pair, cell representations in the “scVI”” latent space were filtered using a 2D PCA Kernel Density Estimation to remove low-density outliers below the 1^*st*^ percentile. For donors meeting the minimum threshold of 50 cells, 500 cells were sampled with replacement to standardize sample size across donors. Sub-clustering was then performed on the sampled “scVI” representations using the Leiden algorithm (*k* = 30 nearest neighbors, resolution = 0.8). Raw UMI counts within each cluster were aggregated to generate sub-donor pseudo-bulk expression profiles. In both datasets, generating log(CPTT + 1) with the pseudo-bulk counts is straightforward. Pseudo-bulk counts were scaled relative to the total sequencing depth of sample *j* to a target sum of 10^4^:

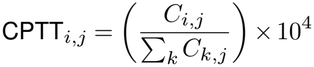

where the denominator represents the sum of raw aggregated counts across all genes *k* in pseudo-bulk sample *j*. In the SN Hormone dataset, log_2_ FC_*ij*_ was possible and was conducted at the pseudo-bulk level in the same way as outlined above for PTI and Hormone datasets.

The Leaf dataset (Kim et al., 2021) was directly transformed from raw reads into log(CPTT + 1), as outlined above. Finally, the PBMC dataset (Zheng et al., 2017) were processed using Scanpy (Wolf et al., 2018). Cells were filtered to retain those with 200 *< N*_genes_ *<* 4000 and mitochondrial read fraction *<* 15%, while genes expressed in fewer than 3 cells were excluded. Raw counts were normalized to log(CPTT + 1), as outlined above.

### Structured train-test split, hyperparameter tuning and model fitting

To prevent data leakage arising from gene homology, both the outer 90*/*10 train-test split and the inner CV folds were structured with respect to target gene families (Washburn et al., 2019; Li et al., 2024). This partitioning guarantees that all target genes in the same family are assigned exclusively to either the training or evaluation set. Gene families were derived by performing an all-vs-all protein sequence alignment using BLASTP, retaining significant homologous pairs (*E*-value *<* 10^−5^, bitscore *>* 40). From these alignments, an undirected network was constructed with bitscores as edge weights and partitioned into distinct clusters via the Markov Clustering (MCL) (Van Dongen, 2008) algorithm with an inflation factor of 1.1. Any genes lacking significant alignments were assigned to unique single-member families to ensure complete genome coverage.

Model hyperparameters were optimized over 50 trials using the Optuna framework (Akiba et al., 2019) with a multivariate Tree-structured Parzen Estimator (TPE) sampler. The hyperparameter search spaces were configured according to the default parameters: Overall, penalty strength *λ* was sampled logarithmically over *λ* ∈ [10^−4^, 2.0], Elastic Net mixing ratio *α* was sampled uniformly over *α* ∈ [0, 1.0]. To allow rapid exploration, single precision (GPU is faster) was used during the CV runs, with an *ϵ* (see Algorithm 1) of 0.001 with a maximum of 5000 iterations. Note that tight convergence is difficult to achieve with single precision, hence the relatively high tolerance and low iterations.

Each trial executed a 3-fold structured CV scheme within the training set. Candidate hyperparameter sets were evaluated by maximizing the average out-of-fold Pearson correlation coefficient between observed gene expression and predicted expression across all folds. After the optimal hyperparameters are obtained, a new model is optimized on the full training data with the selected *λ* and *α*. Performance is evaluated on the held-out test set. Finally, a new model is trained over all (training and test) data. That model is used for downstream tasks such as TFBM inference (see Figure S12). For these models, double precision with *ϵ* = 1 · 10^−8^ and a maximum of 200.000 iterations. All hyperparameter tuning and fitting was conducted on an H100 GPU.

### Importance score calculation

Let us refer as *H* to the Hessian matrix of the smooth part of ref Equation 7 (see Equation 8). To quantify the relevance of the inferred TF-motif interactions, we define an importance score based on the magnitude of the coefficient:

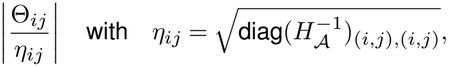

where *H*_A_ denotes the principal submatrix formed by selecting rows and columns of *H* indexed by A, the set of indices of non-zero coefficients in Θ. Note that diag(*H*_A_) must be calculated directly, leveraging the sparsity of Θ, since *H* cannot be computed explicitly due to its memory requirements. This attempts to refine the raw |Θ_*ij*_ | coefficient by accounting for local geometry of the objective around the fitted solution, ensuring that interactions are ranked by the plausibility of the interaction rather than relying solely on the raw magnitude |Θ_*ij*_ |.

### Motif reconstruction

To translate the inferred *k*-mer model weights into biologically interpretable TFBMs, we developed an iterative anchor-based alignment strategy to construct PPMs directly from the importance scores assigned to each parameter value in Θ. For a given TF *j*, the set of active *k*-mers was extracted. The *k*-mer exhibiting the maximum importance, argmax 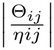, was designated as the anchor sequence.

All other active *k*-mers associated with the TF were aligned to this anchor using a global pairwise alignment algorithm. To account for the bidirectional nature of DNA binding, both the forward and reverse complement sequences of each *k*-mer were evaluated, and the orientation yielding the maximum alignment score was retained. Alignment parameters were strictly configured to prohibit internal and discourage pointless terminal gaps (scores of −100 and −0.2, respectively), thereby preventing the artificial fragmentation of *k*-mers and penalizing trivial, non-overlapping alignments. Furthermore, *k*-mers with an optimal alignment score falling below 25% of the anchor’s self-alignment score were discarded as noise.

Following alignment, *k*-mers were mapped to a unified positional profile relative to the anchor. At each position *k* and for each nucleotide *b* ∈ {*A, C, G, T* }, a weighted frequency was calculated by summing importance scores of all aligned *k*-mers containing nucleotide *b* at that position. This weighted profile was subsequently row-normalized to yield a formal PPM.

To ensure the biological robustness and specificity of the reconstructed motifs, we calculated the total Information Content (IC) for each PPM:

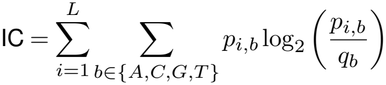

where *L* is the length of the aligned profile, *p*_*k,b*_ is the probability of nucleotide *b* at position *k*, and *q*_*b*_ = 0.25 represents a uniform background nucleotide distribution. Reconstructed motifs with a total IC below a threshold of 10 bits (a unique 6-mer has an IC of 12) were deemed low-confidence and excluded from further analysis. The final, filtered, set of motifs was exported in standard MEME format to facilitate downstream motif comparison and annotation.

### JASPAR comparison and QQ-Plot construction

To evaluate the biological validity of the inferred motifs, we systematically compared our reconstructed PPMs against empirically validated binding profiles from reference databases, in particular JASPAR 2026 (Ovek Baydar et al., 2026) non-redundant MEME-format plant data for *Arabidopsis thaliana* and the analogous version for vertebrates for human. Motif alignment and similarity scoring were performed using the TOMTOM algorithm from the MEME Suite (Gupta et al., 2007). The high-confidence inferred motifs (exported in MEME format) served as queries against the non-redundant JASPAR databases. TOMTOM was executed with a maximally permissive initial significance threshold (-thresh 1) to retain all the comparisons in the final file.

To assess whether the observed TOMTOM matches exceeded what would be expected by chance, we constructed QQ-plots of the observed −log_10_ p-values distribution against the expected distribution under the null hypothesis of uniformly distributed p-values. For each dataset independently, p-values were ranked in ascending order and the *i*^th^ ranked p-value was assigned an expected p-value of *i/*(*n* +1), where *n* is the total number of TOMTOM comparisons for that dataset (equal to the number of TFs for Figure 3a and considerably larger for panel b). Both observed and expected p-values were −log_10_ transformed and plotted against each other, with the identity line (*y* = *x*) representing the null expectation of no enrichment. Deviation above the diagonal reflects an excess of low p-values relative to chance, indicating enrichment of true motif similarity beyond background. For the top-ranked hit in each dataset, the corresponding query motif consensus sequence and its JASPAR match were annotated directly on the plot.

## CODE AVAILABILITY

Code to use BRIDGE and reproduce this paper is available at https://github.com/JordiAlonsoEsteve/BRIDGE.

## ACKNOWLEDGEMENTS

This publication is part of the long-term program PlantXR: A new generation of breeding tools for extra-resilient crops (KICH3.LTP.20.005) which is financed by the Dutch Research Council (NWO), the Foundation for Food Agriculture Research (FFAR), companies in the plant breeding and processing industry, and Dutch universities. These parties collaborate in the CropXR Institute (www.cropxr.org) that is funded through the National Growth Fund (NGF) of the Netherlands.

The authors thank Jordan Earle, Julia Ruiz Capella, Anna Neefjes, Samara Almeida Landman, Guido Van den Ackerveken, Huub Hoefsloot, Petra Bleeker, Xenja Ploeger and Rob Schuurink for helpful discussions.

## Supplementary Information

### Standard deviation distributions across datasets

Figures S1 to S7 show the distribution of standard deviations per gene (left panel in each plot) and per sample (right panel in each plot) across the analyzed datasets. As observed across expression units, TPM-normalized datasets consistently exhibit higher standard deviations across samples than across genes, whereas log_2_ FC transformations demonstrate similar variance distributions.

**Figure S1.**
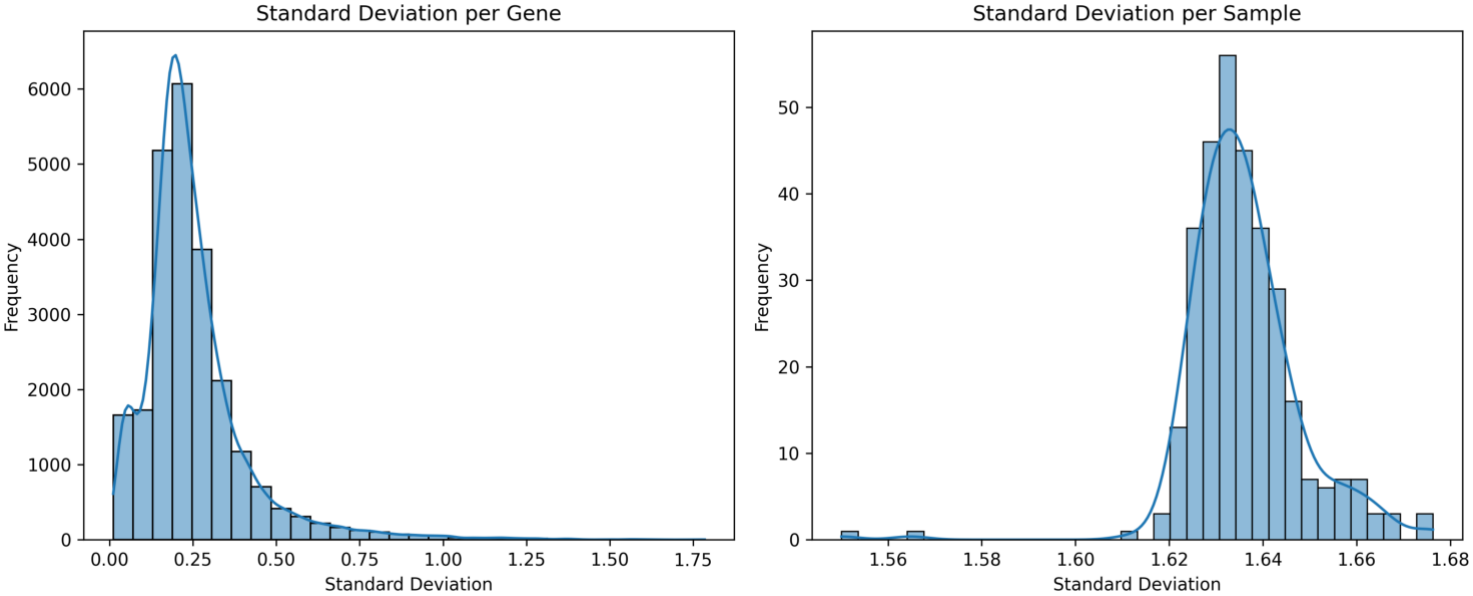
Standard deviation distribution for PTI (TPM)

**Figure S2.**
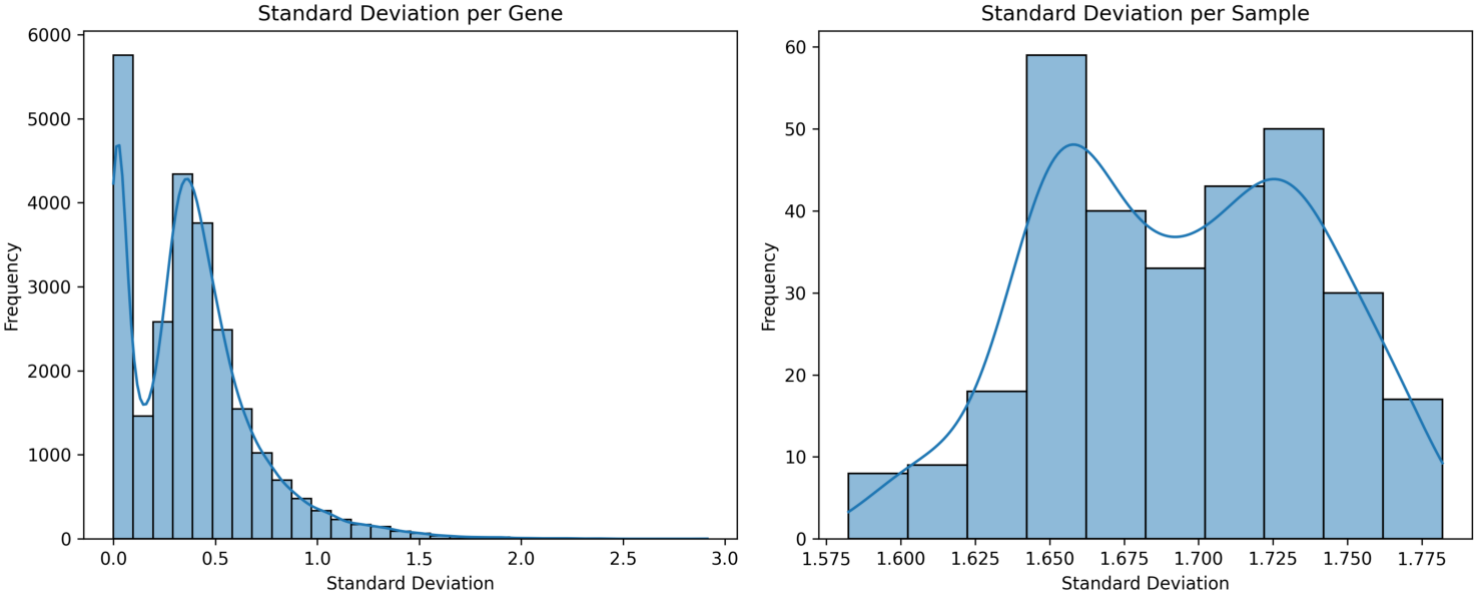
Standard deviation distribution for Hormone (TPM)

**Figure S3.**
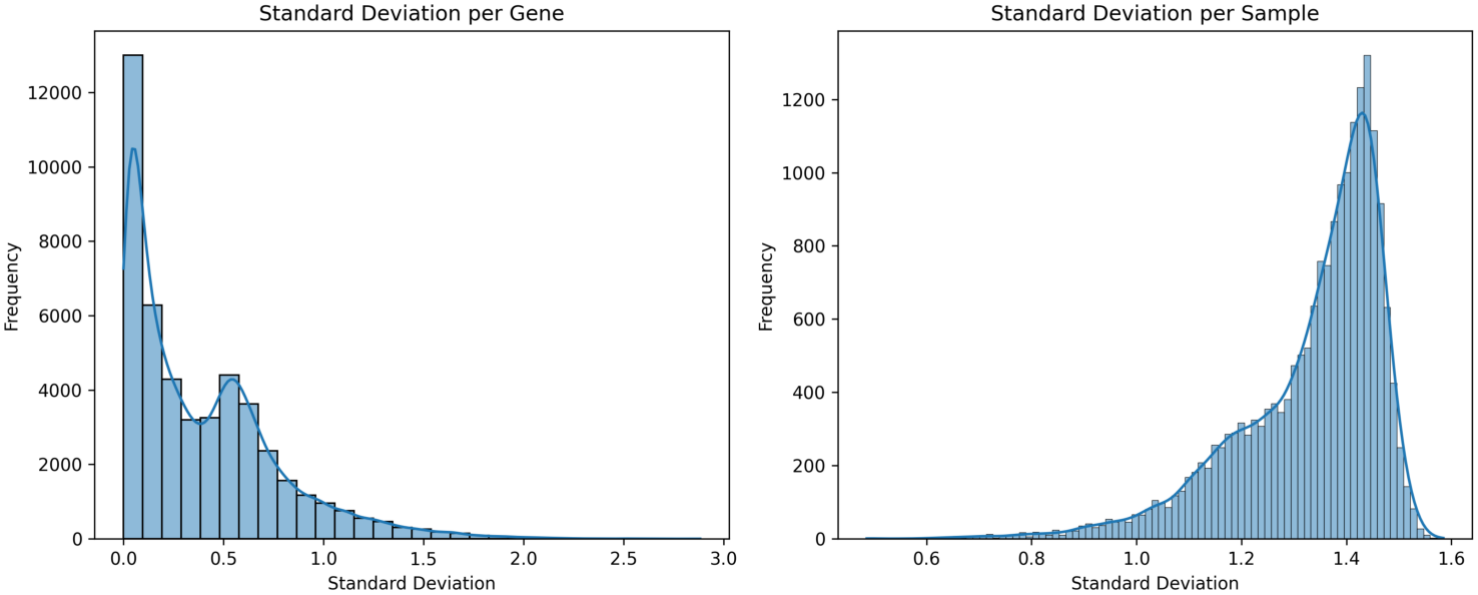
Standard deviation distribution for GTEx (TPM)

**Figure S4.**
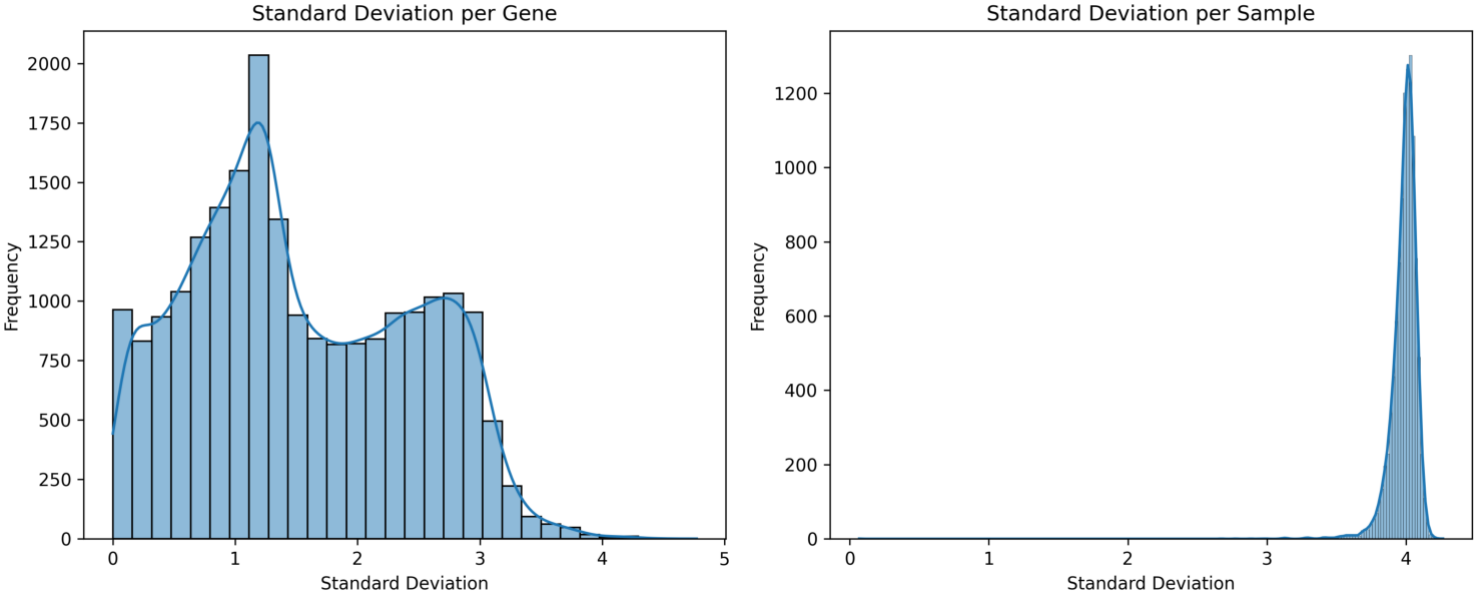
Standard deviation distribution for TCGA (TPM)

**Figure S5.**
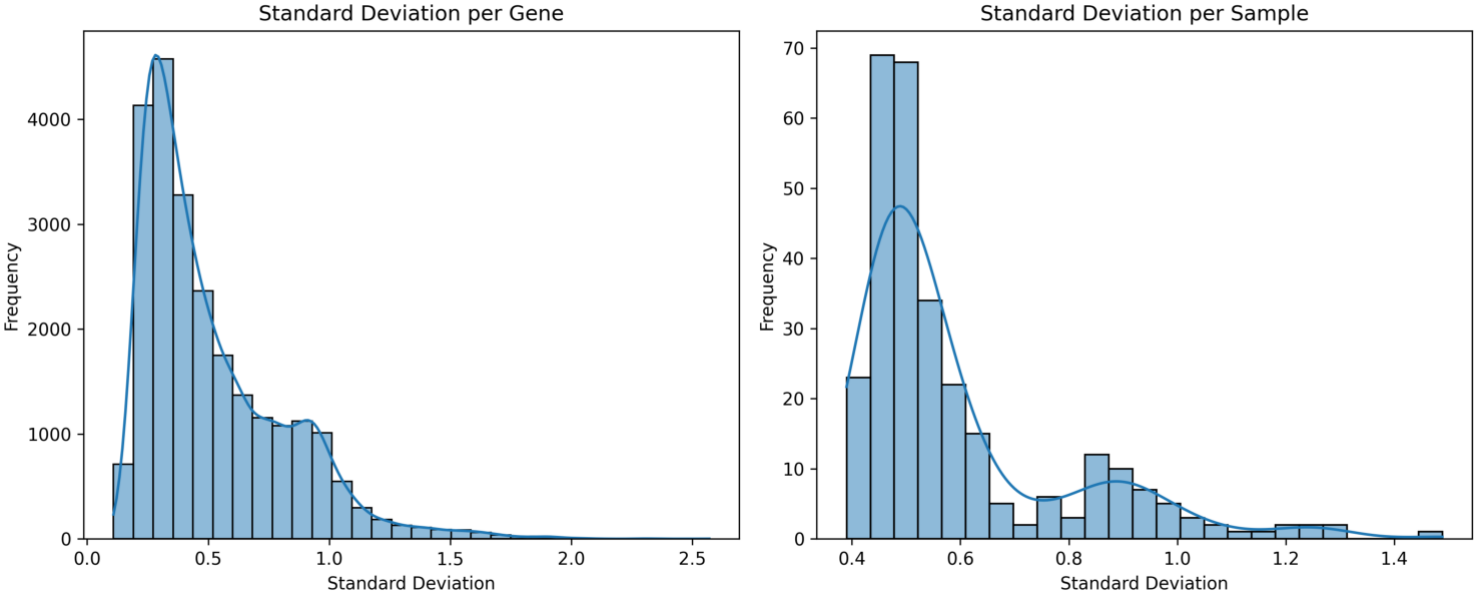
Standard deviation distribution for PTI (log_2_ FC)

**Figure S6.**
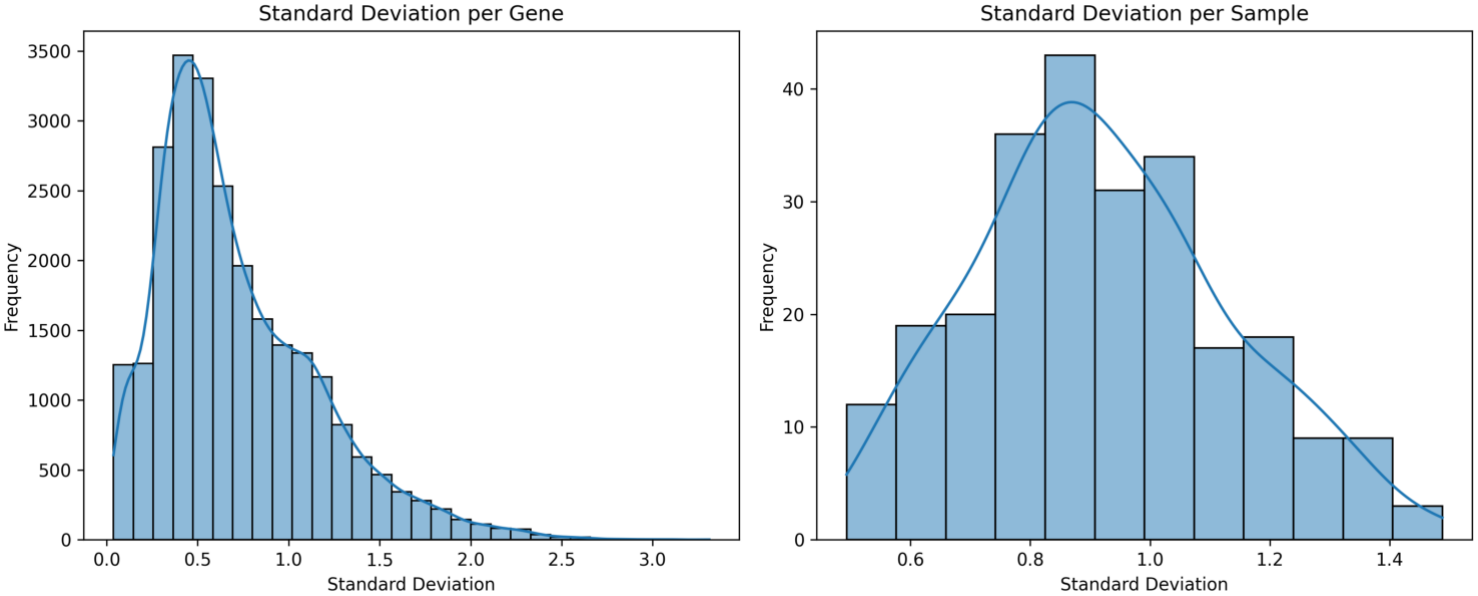
Standard deviation distribution for Hormone (log_2_ FC)

**Figure S7.**
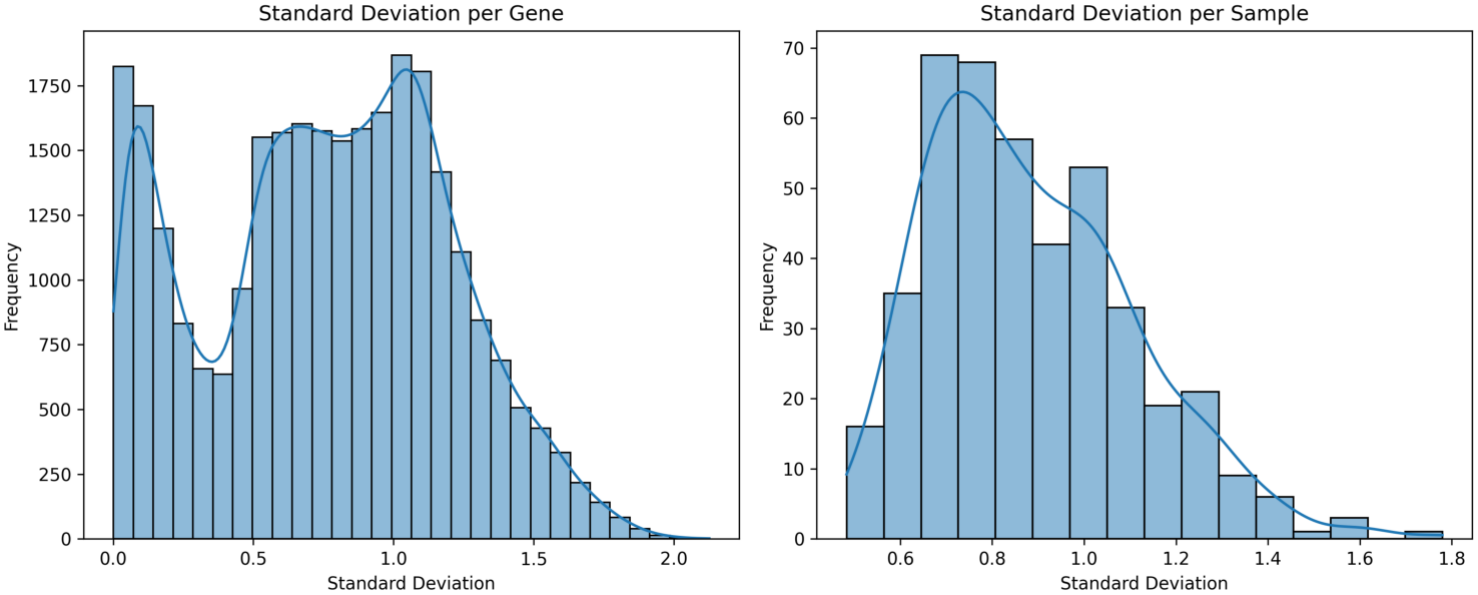
Standard deviation distribution for SN Hormone (log_2_ FC)

**Figure S8.**
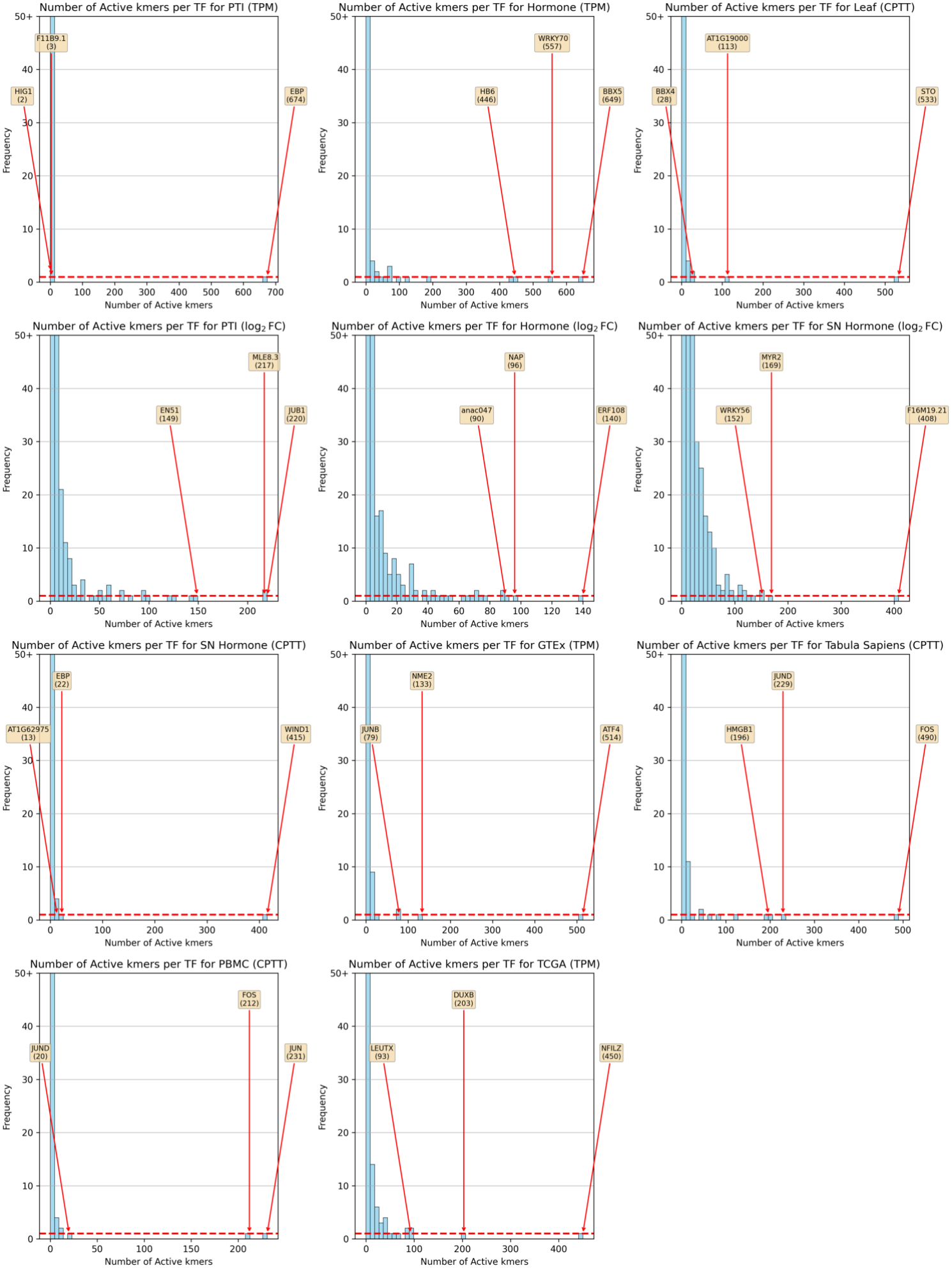
Non-zero 6-mers per TF, distributions across all datasets. Histograms display (using bins of 50) the distribution of the number of non-zero 6-mers (rows) per TF (column) in Θ. This is, the distribution of how many 6-mers are the TFs predicted to bind. On the horizontal axis the number of active 6-mers per TF is displayed and on the vertical axis the number of times that a TF has that number of 6-mers among all the considered TFs. The long tails are particularly problematic because they imply a a TF that binds to many different 6-mers, which contradicts our assumptions regarding binding specificity.

**Figure S9.**
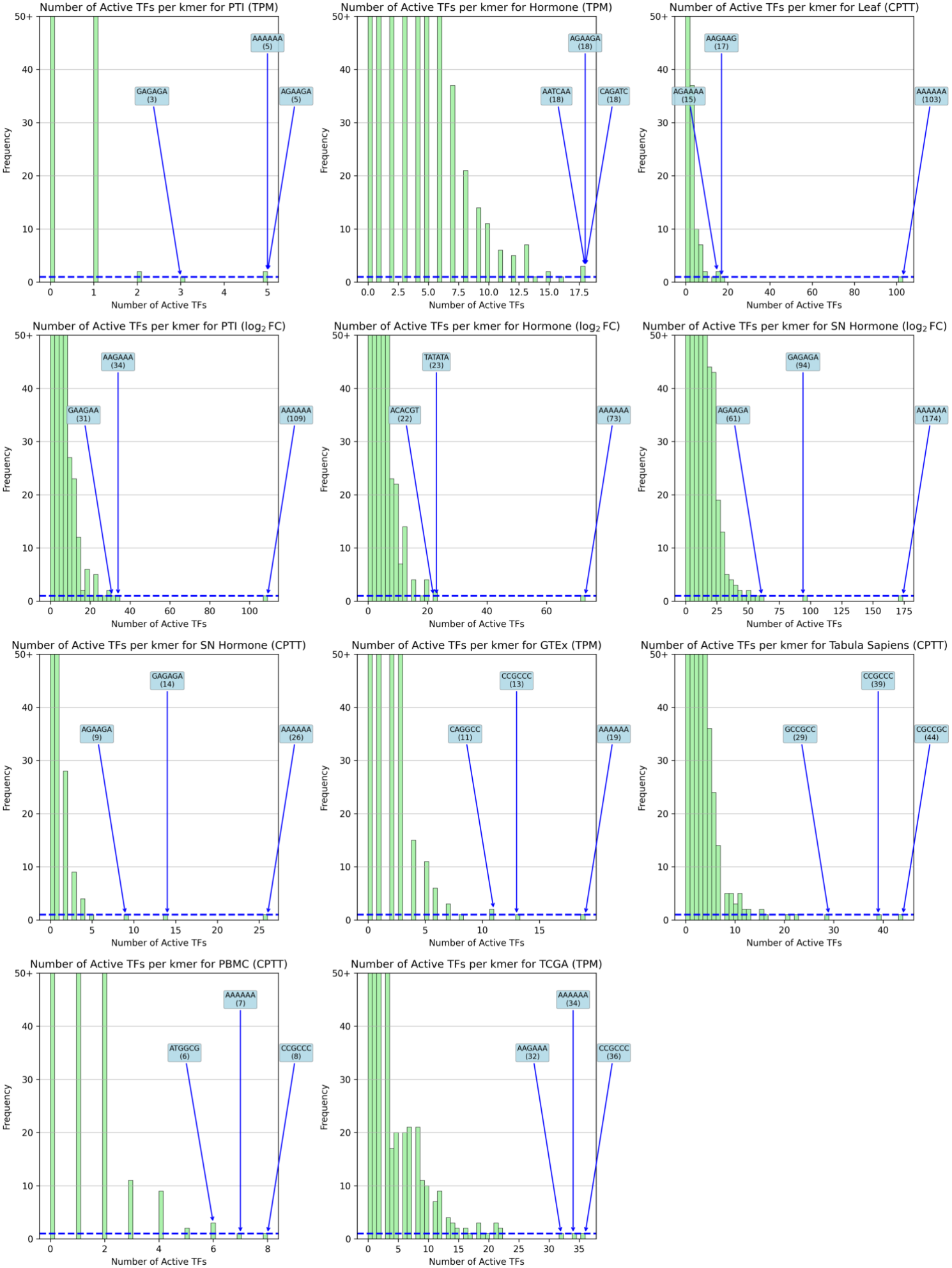
Non-zero TFs per 6-mer, distributions across all datasets. Histograms display (using bins of 50) the distribution of the number of non-zero TFs (colum) per 6-mer (row) in Θ. This is, the distribution of how many TFs are predicted to bind to a particular 6-mer. On the horizontal axis the number of active TF per each 6-mer is displayed and on the vertical axis the number of times that a 6-mer has that level of activity across TFs. As before, long tails are particularly problematic because they imply a 6-mers are active in many TFs, which contradicts the claim of specificity of 6-mers to particular TFs.

**Figure S10.**
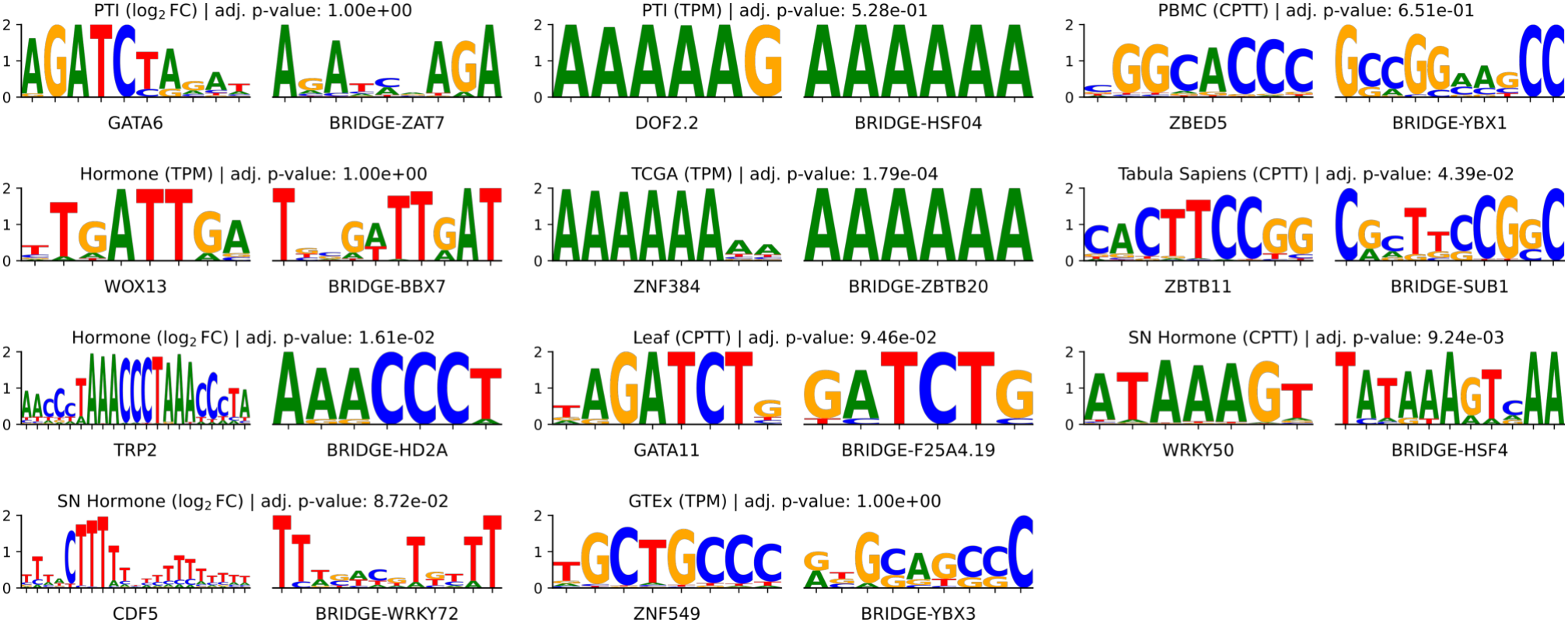
Inferred TFBMs for TF that are not present in the JASPAR database. The most important TFBMs that have an information content above 10. On the left, the most similar motif in the JASPAR database, on the right the BSR inferred motif. Adjusted p-value here refers to the q-value provided by TOMTOM, which controls for all comparisons.

**Figure S11.**
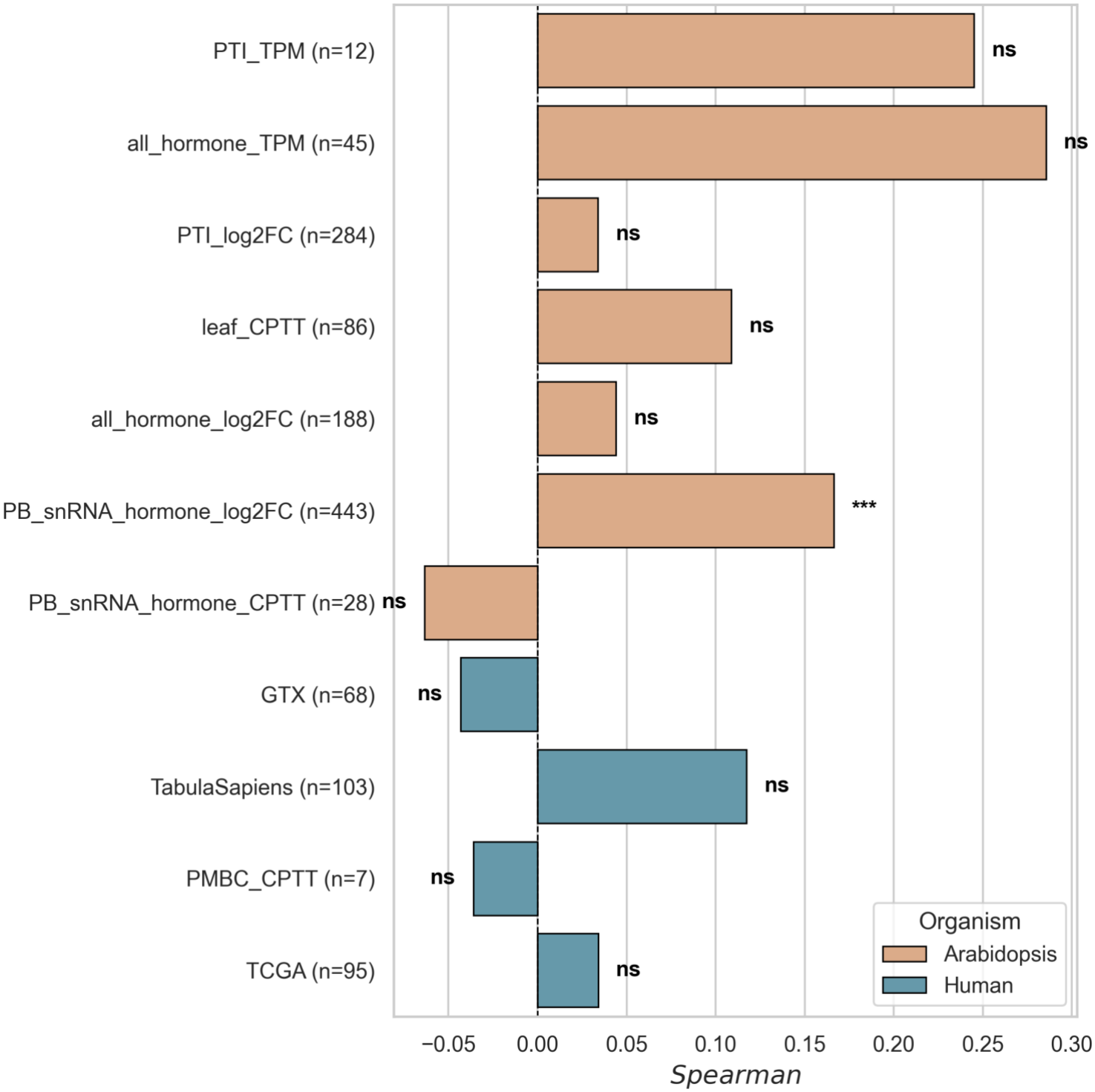
Correlation between TOMTOM’s raw −log_10_ pvalue and importance score of anchor 6-mer with the closest TFBMs within the TF’s family. The number of observations increases because most predicted TFBM now have a counterpart in JASPAR, as most TF families have at least one instance.

**Figure S12.**
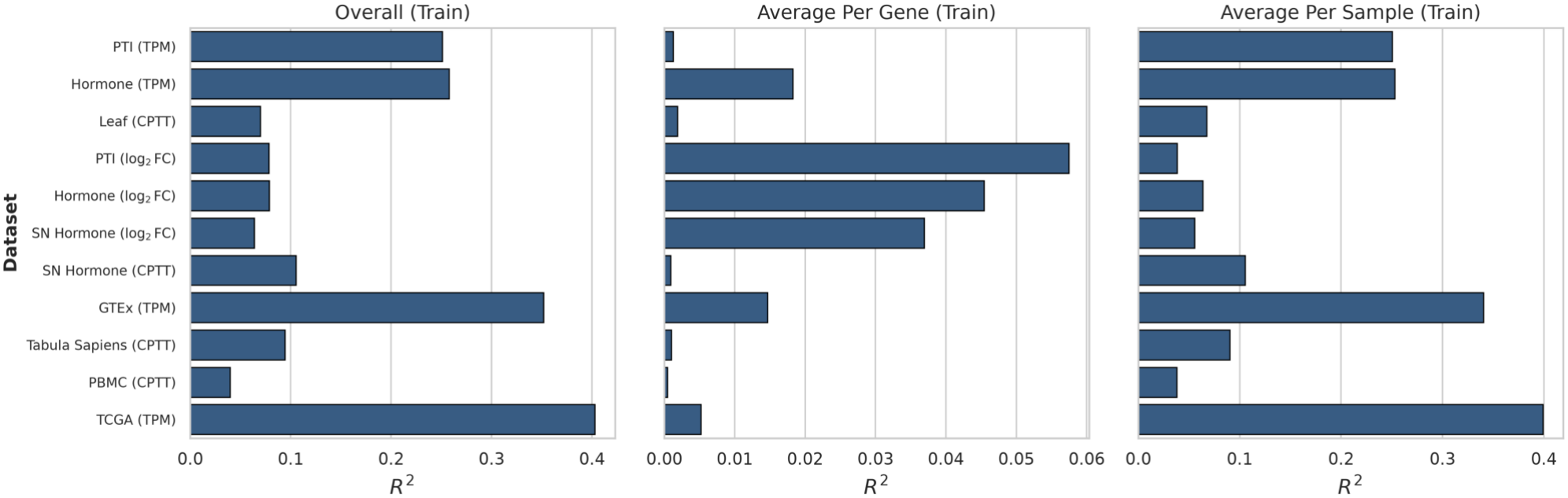
*R*^2^ for the model fitted on the whole data. This is not test set performance. Once the hyperparameter tuning has concluded and we have evaluated the predictive performance in unseen data, the model is re-fitted using all available data for TFBM inference. The *R*^2^ displayed here corresponds to the reconstruction of all training data with this last model.

## Supplementary Tables

**Table S1.**
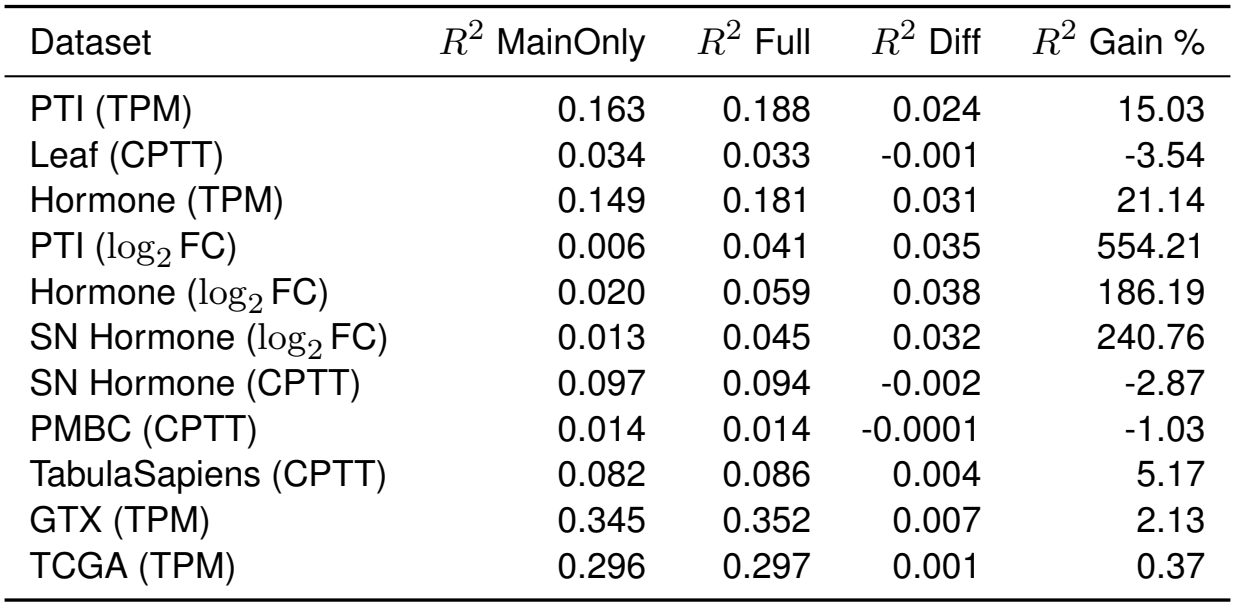
Overall *R*^2^ comparisons breakdown. The inclusion of interaction terms improves predictive performance in most of the datasets.

